# Visual object recognition in the fly: the internal representation of a predator drives defensive behaviour

**DOI:** 10.64898/2026.11.09.765114

**Authors:** Sachira Denagamage, Merrin Foltz, Valentin M. Dalstein, Rudy Behnia, Gwyneth M. Card, Richard Axel, Kevin M. Cury

**Affiliations:** Zuckerman Mind Brain Behavior Institute, Columbia University, New York, NY, USA; Howard Hughes Medical Institute, Columbia University, New York, NY, USA

## Abstract

Animals recognize visual objects and respond to them appropriately to survive. This process is thought to unfold in stages: the visual scene is first deconstructed into constituent features, which are then reconstructed into representations of object identity and relayed to motor circuits. Here we define a neural circuit in *Drosophila* that underlies the visual recognition of a natural predator, the parasitoid wasp, and orchestrates defence. We found that flies exhibit a repertoire of directed defensive actions structured by the wasp’s position within the fly’s field of view. Expression of these behaviours is driven by visual detection of a conspicuous morphological cue — the wasp’s elongated antennae. A small subset of lobula columnar (LC) visual projection neuron types are required for defence and are activated by constituent visual features of the wasp. Three LC types respond selectively to the wasp’s antennae, and relay this information to LC9, which sums antennal- and body-feature signals to form a composite representation of the wasp. Spatial filtering of this representation onto a descending premotor pathway translates the wasp’s location in visual space into directed defence. Feature combination followed by spatial filtering may constitute a general mechanism for coupling object identity to context-appropriate action.

## Main

Organisms have evolved mechanisms to extract visual information from the environment and process this information to create an internal representation of the external world. These representations support the recognition of objects and guide context-appropriate behaviours essential for survival. Visual recognition of other animals, including potential mates, rivals, and predators, drives behaviours ranging from courtship and aggression to vigilance and evasion. For insects, parasitoid wasps are a widespread predatory threat — female wasps lay eggs in host larvae, pupae, or adults, resulting in death as the wasp offspring develop ^1,2^. In *Drosophila*, vision is critical for detecting and responding to parasitoid wasps^3–5^; however, the visual cues that enable this recognition have not been identified.

Appropriate responses to natural objects often require that the nervous system deconstruct objects into constituent visual features and then reconstruct them into a more complete representation that is conveyed to motor circuits. In the fly, the visual world is deconstructed into features that are encoded by lobula columnar neurons (LCs), visual projection neurons that carry information from the optic lobe to the central brain and comprise approximately 50 distinct cell types ^6–9^. Each LC type responds to specific features of simple visual stimuli, such as the size of small moving dots or thin bars, or the expansion rate of looming disks ^10,11^. Activation of some LC types is sufficient to evoke behaviours such as backwards walking or takeoff ^7^ — a feature-to-action mapping that is reminiscent of the vertebrate tectum and superior colliculus ^12–14^. Natural objects such as parasitoid wasps, however, comprise multiple visual features distributed across distinct morphological components, raising the question of whether and how signals across LC pathways are combined to reconstruct a more complete representation that guides behaviour.

Object recognition has been studied extensively in the primate ventral stream, a visual processing pathway where object-level representations are thought to arise hierarchically through progressive reconstruction ^15,16^. Successive stages support increasingly complex representations, from orientation to contours and textures, and ultimately, to complex objects such as faces ^17–21^. In the inferior temporal cortex, neurons responsive to select facial features are presumed to converge to reconstruct a more complete facial image ^22–24^. However, the circuitry through which features are combined to represent complete objects remains largely inaccessible. Moreover, because studies of the ventral stream have focused primarily on perception, the output of the system has not been linked to behaviour.

Defence against a natural predator in *Drosophila* offers an opportunity to trace how a natural object is deconstructed into visual features, reconstructed into an object-level representation, and transformed into action within a synaptically resolved nervous system. In this study, we identify wasps’ elongated antennae as a key visual cue driving acute defence in flies, and describe a circuit that integrates visual features of distinct morphological components of a wasp into an object-level representation, then spatially filters this representation to generate directed, defensive action.

### Vision mediates evasion of a natural predator

We examined the behavioural response of adult fruit flies to parasitoid wasps by video tracking individual female flies ^25^ (*Drosophila melanogaster*) paired with a male or female wasp (*Leptopilina heterotoma*) in a small rectangular chamber (**Supplementary Videos 1–3**; **Fig. 1a**). As a control, we paired flies with a different drosophilid species. Flies were more active when paired with wasps than with control flies, as measured by chamber midline-crossing rate (**Extended Data Fig. 1a**). Fly and wasp velocities were highly correlated, with increases in male wasp velocity preceding increases in fly velocity by 1.7 s on average (**Fig. 1b**), indicating that wasp movement induced fly movement. We also identified three evasive fly behaviours observed in the fly–wasp pairings — forward running, backing up, and jumping (**Fig. 1c, Extended Data Fig. 1b**; **Supplementary Videos 4–6**) — all of which acutely increased the inter-animal distance (**Extended Data Fig. 1c**). Jumps often occurred after contact (52% occurring within 50 ms following contact), compared to only 7% and 10% for forward running and backing up, respectively (**Fig. 1c**). Consistent with these evasive reactions, average inter-animal distances over the entire interaction were significantly greater for fly–wasp pairings than for fly–fly pairings (**Fig. 1d**). Together, these data indicate that adult flies actively evade parasitoid wasps, using directed locomotion to maintain distance and jumping when contact occurs. Interestingly, we noted that male wasps elicited stronger evasion than females even though the females are predatory and the males pose no threat (see Discussion). We therefore used male wasps in subsequent experiments, except where noted.

**Figure 1.**
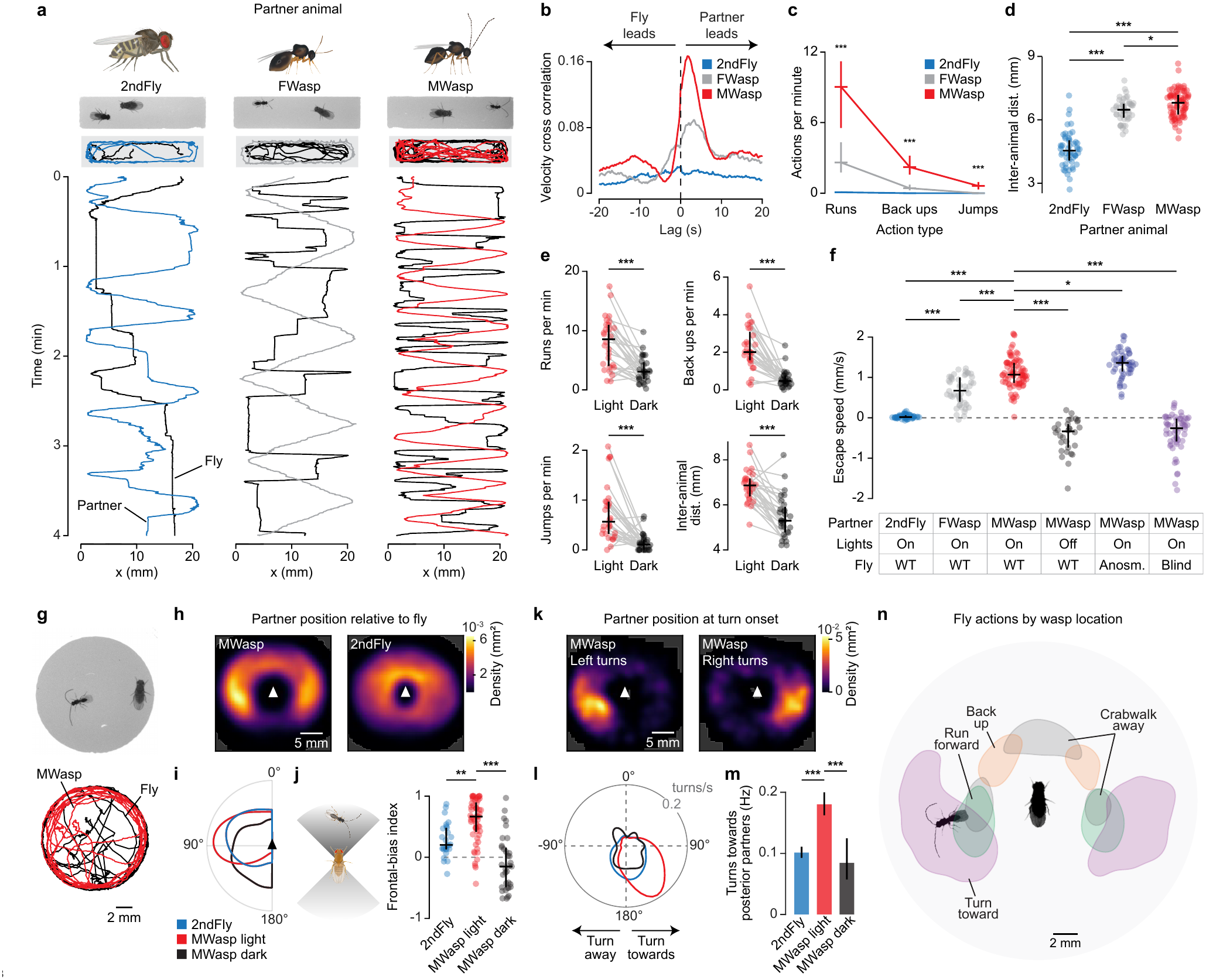
Visually mediated evasion and monitoring of a natural predator. **a,** Representative examples of rectangular two-animal interaction assay. Top image: screenshot of assay. Top traces: x-y position of each animal’s central midpoint over the initial 2 min of the lights-on session. Bottom traces: x-position over the initial 4 min. A female *D. melanogaster* fly (black traces) paired with a male heterospecific fly (left, blue traces; 2ndFly; *D. micromelanica*), a female wasp (middle, grey traces; FWasp; *L. heterotoma*), or a male wasp (right, red traces; MWasp; *L. heterotoma*). Corresponding videos are in Supplementary Videos 1–3. **b,** Mean normalized cross-correlation of the velocity of the two animals in each pairing type. Here and in c and d, *n* (pairs): 2ndFly, 52; FWasp, 48; MWasp, 82. Positive lag indicates that fly velocity lags partner velocity. **c,** Rate of forward-running, backing-up, and jumping events performed by flies paired with three distinct partner animals. Horizontal lines, median; vertical lines, interquartile range (25^th^–75^th^ percentile). One-way ANOVAs (all df = 2, 179): Forward run, *F* = 148.51, *P* = 9.61 × 10^−39^; Back up, *F* = 159.36, *P* = 1.78 × 10^−40^; Jump, *F* = 84.07, *P* = 1.80 × 10^−26^. Tukey–Kramer post hoc tests: Forward run, 2ndFly vs. FWasp, *P* = 5.49 × 10^−8^, and 2ndFly vs. MWasp, *P* = 5.47 × 10^−38^; Back up, 2ndFly vs. FWasp, *P* = 2.28 × 10^−3^, and 2ndFly vs. MWasp, *P* = 1.84 × 10^−37^; Jump, 2ndFly vs. FWasp, *P* = 0.701, and 2ndFly vs. MWasp, *P* = 2.99 × 10^−22^. **d,** Average inter-animal distance across three pairing types. Here and in e, f, j: points, individual pairings; horizontal lines, median; vertical lines, interquartile range (25^th^–75^th^ percentile). One-way ANOVA: *F* (2, 179) = 178.62, *P* = 2.25 × 10^−43^. Tukey–Kramer post hoc test: 2ndFly vs. FWasp, *P* = 5.39 × 10^−64^; 2ndFly vs. MWasp, *P* = 7.78 × 10^−82^; FWasp vs. MWasp, *P* = 0.032. **e,** Rate of forward-running, backing-up, and jumping events, as well as the inter-animal distance, for flies paired with male wasps, measured in paired lights-on and dark sessions. *n* = 28 pairs. Paired t-test on light versus dark: Forward run, *P* = 2.38 × 10^−6^; Back up, *P* = 7.16 × 10^−8^; Jump, *P* = 1.98 × 10^−5^; inter-animal distance, *P* = 1.38 × 10^−7^. Wasp activity was similar in both lights-on and dark sessions (Extended Data Fig. 1d). **f,** Escape speed of flies across partner and lighting conditions. *n* (pairs): 2ndFly, 52; FWasp, 48; MWasp-light, 82; MWasp-dark, 30; Anosmic, 51; Blind, 57. One-way ANOVA: *F* (5, 314) = 208.50, *P* = 1.71 × 10^−97^. Tukey–Kramer post hoc test: 2ndFly vs. FWasp, *P* = 2.16 × 10^−16^, and MWasp-light, *P* = 1.60 × 10^−35^; MWasp-light vs. FWasp, *P* = 4.52 × 10^−9^, MWasp-dark, *P* = 1.52 × 10^−44^, Anosmic, *P* = 0.0200, and Blind, *P* = 6.63 × 10^−55^. **g,** Top: Screenshot from representative example of circular paired-animal interaction assay for fly paired with male wasp. Bottom: x-y position of each animal’s central midpoint over the initial 2 min. **h,** Probability density of partner-animal position relative to the fly for pairings with male wasps (left; *n* = 57 pairs) and male flies of a second species (right; *n* = 26 pairs). Central arrowhead depicts fly position with the fly facing upwards. Only frames occurring while the fly was on the chamber floor or ceiling were considered. Here and in k: grey masking indicates positions for which the density was 0. **i,** Kernel density estimate (a.u.) of partner-animal azimuthal position relative to the fly, using a Gaussian kernel (bandwidth = 10°); right hemisphere data was mirrored about the 0°–180° azimuthal axis and added to left hemisphere data. Same data as in h. **j,** Frontal-bias index across partner and lighting conditions. *n* (pairs): 2ndFly, 26; MWasp-light, 57; MWasp-dark, 35. One-way ANOVA: *F* (2, 115) = 38.41, *P* = 1.67 × 10^−13^. Tukey–Kramer post hoc test: MWasp-light vs. 2ndFly, *P* = 1.83 × 10^−3^; MWasp-light vs. MWasp-dark, *P* = 6.00 × 10^−14^. **k,** Probability density of male wasp position relative to the fly at the onset of fly leftward turns or fly rightward turns. *n* (turns; from 57 pairs): leftward, 819; rightward, 755. Central arrowhead depicts fly position with the fly facing upwards. Only events with onsets more than 5 s following inter-animal contact were considered here. **l,** Turning rate as a function of partner’s relative azimuthal position. *n* (turns, pairs): 2ndFly, 2159, 26; MWasp-light, 1574, 57; MWasp-dark, 100, 35. Partner position was mirrored about the 0°–180°azimuthal axis for left turns such that all turns towards the wasp (ipsiversive turns) are depicted on the right hemisphere of the plot. **m,** Rate of turning towards the partner animal (ipsiversive turns) for turns initiated when the partner animal was located in the lateral-posterior visual field, between 121°and 177°azimuth, on the same side as the turn (see Methods). Error bars, Wilson 95% confidence intervals. *χ*^2^ tests on pooled counts, Holm-corrected across all three pairwise comparisons: MWasp light vs. 2ndFly, *P* = 3.33 × 10^−16^; MWasp dark vs. 2ndFly, *P* = 0.376; MWasp dark vs. MWasp light, *P* = 3.49 × 10^−4^. *n* turns: 2ndFly, 475; MWasp light, 353; MWasp dark, 25. **n,** Iso-contour maps of smoothed event counts of male wasp position relative to the fly at the onset of distinct behavioural actions. *n* (events; from 57 pairs): Forward run, 3150; Back up, 94; Turn, 1574; Crabwalk, 981. Contours drawn at 50% of the peak of each distribution. Only events occurring while the fly was on the chamber floor and with onsets more than 5 s following inter-animal contact were considered here.

We determined whether fly evasive reactions were visually mediated by analysing fly–wasp pairings both in the dark and in genetically blind *norpA* flies with blocked phototransduction. Fly evasive behaviours were nearly abolished in darkness (**Fig. 1e, Extended Data Fig. 1d**), and the few remaining bouts of forward running and backing up were primarily contact triggered (**Extended Data Fig. 1e**). Average fly–wasp distance was also significantly reduced in the dark (**Fig. 1e**). We defined a metric, escape speed, to quantify how rapidly flies moved away from the animal they were paired with during close, non-contact encounters (see Methods). Flies in the dark and genetically blind flies exhibited a remarkably diminished escape speed, near zero, indicating that non-contact evasive responses were abolished (**Fig. 1f**). In contrast, flies in which olfactory and auditory input were removed by excising the antennae and maxillary palps escaped with speeds comparable to wild type flies in the light (**Fig. 1f**). Together, these data show that flies rely on vision, and not audition or olfaction, to recognize parasitoid wasps as a possible threat and execute evasive manoeuvres before contact.

### Predator location drives directed turning and evasive manoeuvres

Flies may also visually monitor the wasp, an anti-predator vigilance behaviour that reduces the risk of ambush. We conducted a new set of experiments in a circular chamber with a larger base area, enabling freer movement and more precise quantification of the angular relationship between animals (**Fig. 1g**; **Supplementary Video 7**). We found that the wasp was frequently positioned to the front and side of the fly and was rarely found behind the fly, in its blind spot ^26^ (**Fig. 1h, i**). We calculated a frontal-bias index comparing partner occupancy in the fly’s anterior versus posterior visual field (see Methods). The index approached +1 for male wasp pairings in the light, indicating the wasp was positioned anterior to the fly, and the index was significantly reduced in darkness or when flies were paired with heterospecific control flies (**Fig. 1j**).

We asked whether this frontal bias was accompanied by active reorientation by the fly towards the wasp. We examined the wasp’s position relative to the fly at the onset of individual turns. Flies made directed turns towards the wasp when the wasp was located in the lateral-posterior field (**Fig. 1k**; **Supplementary Video 8**). Such spatially structured turning was minimal in darkness and in pairings with control flies (**Fig. 1l, m**). Together these observations indicate that flies execute directed turns to maintain the wasp within their anterior-lateral visual field.

Other defensive behaviours might likewise be organized by the wasp’s position in the fly’s visual field. We examined three directions of locomotor evasion: forward, lateral (“crabwalks”), and backwards. We computed the distribution of wasp locations relative to the fly at the onset of each behaviour (**Fig. 1n**). Flies initiated forward running when wasps occupied their lateral-posterior visual field, crabwalks when wasps were directly in front or to the side, and backing up when wasps were positioned anterior-laterally. Thus, flies exhibit a stereotyped repertoire of anti-predator behaviours, including evasive and vigilance-related actions, that depend on the position of the wasp within the fly’s field of view.

### Elongated antennae are a key visual cue eliciting fly defence

Which visual cues associated with wasps elicit defensive behaviour in flies? We initially explored this question by exploiting natural variation in morphology and behaviour among males and females of two additional wasp species that parasitize *Drosophila* (**Fig. 2a**, **Extended Data Fig. 2a, b**). We fit linear models to determine which wasp characteristic best predicted variation in two defensive metrics, escape speed and frontal-bias index. We considered six morphological and behavioural characteristics of the wasps — antennal length, body length, time in chamber, midline-crossing rate, speed, and antennal tip velocity. The models explained 76% and 64% of the variance across group means in escape speed and frontal-bias index, respectively. Antennal length emerged as the strongest predictor of both metrics (**Fig. 2b**, **Extended Data Fig. 2c**).

**Figure 2.**
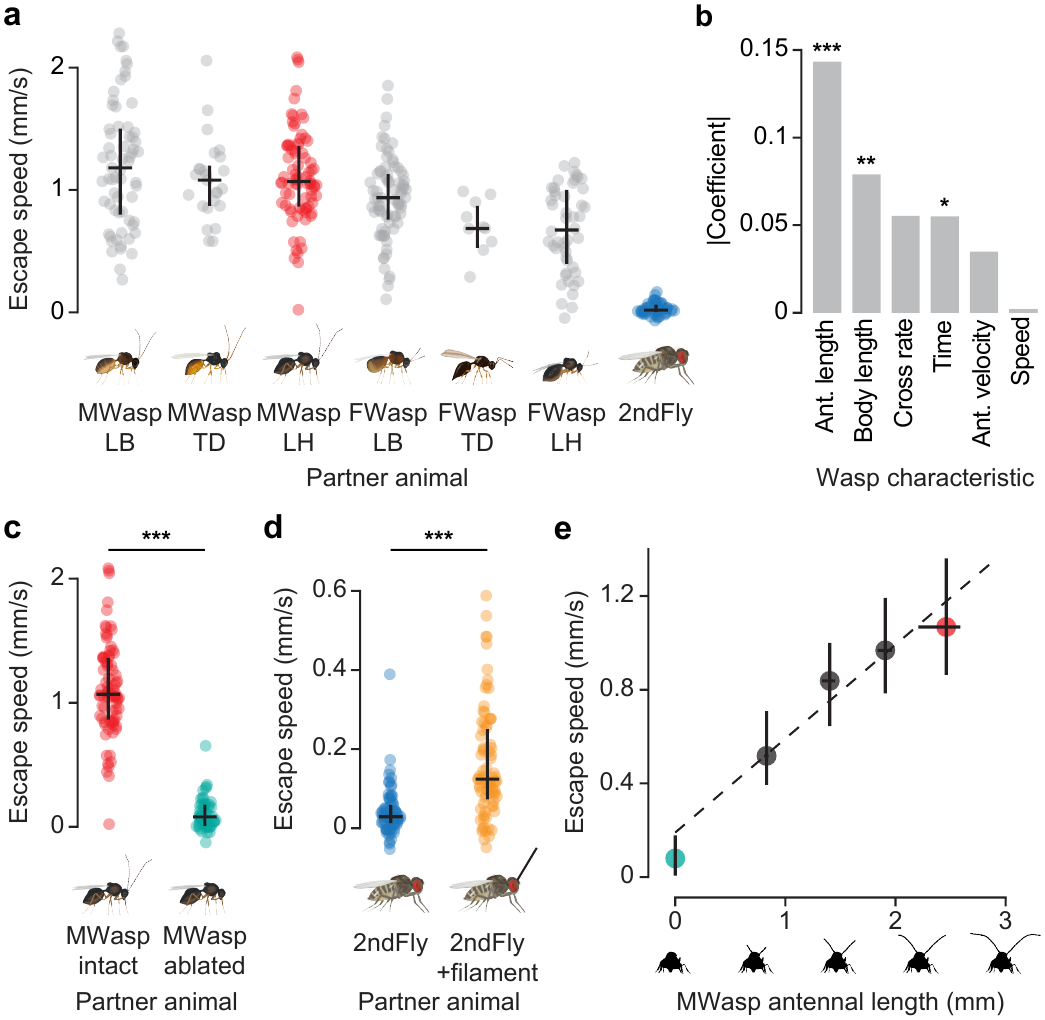
Elongated antennae of wasps are a key visual cue driving fly defence. **a,** Escape speed of flies paired with males or females of three different *Drosophila*-targeting parasitoid wasp species and a male fly of a different species. LB, *Leptopilina boulardi*; TD, *Trichopria drosophilae*; LH, *Leptopilina heterotoma*. Here and in b, *n* (pairs): male *L. boulardi*, 65; male *T. drosophilae*, 25; male *L. heterotoma*, 82; female *L. boulardi*, 80; female *T. drosophilae*, 11; female *L. heterotoma*, 48; male *D. micromelanica*, 52. Here and in c, d: points, individual pairings; horizontal lines, median; vertical line, interquartile range (25^th^–75^th^ percentile). **b,** Coefficient magnitudes from a multiple linear regression model with six morphological and behavioural predictors of fly escape speed. Antennal length, *P* = 1.17 × 10^−7^; body length, *P* = 0.00121; midline-crossing rate, *P* = 0.960; time in chamber, *P* = 0.0319; antennal velocity, *P* = 0.314; speed at midline crossing, *P* = 0.0897. Antennal length, speed at midline crossing, and antennal velocity had positive coefficients; all others, negative. The model explained 76% of the variance in escape speed across group means. **c,** Escape speed of flies paired with individual male wasps (*L. heterotoma*) with intact (MWasp intact) or fully removed antennae (MWasp ablated). *n* (pairs): MWasp intact, 82; MWasp ablated, 50. Two-sided two-sample t-test: *P* = 1.13 × 10^−37^. Removing the antennae had modest to no effects on wasp behaviour (Extended Data Fig. 2g). **d,** Escape speed of flies paired with individual heterospecific flies without (2ndFly) or with (2ndFly+filament) a synthetic filament affixed to their heads. *n* (pairs): 2ndFly, 75; 2ndFly+filament, 80. Two-sided two-sample t-test: *P* = 3.00 × 10^−12^. **e,** Escape speed of flies versus wasp antennal length for pairings with male wasps whose antennae were cut to four distinct lengths, or left intact. *n* (pairs): 0 segments, 50; 3 segments, 61; 6 segments, 60; 9 segments, 58; 13 segments (intact), 82. Linear regression: intercept = 0.190, slope = 0.401, *R*^2^ = 0.545, *P* = 8.53 × 10^−55^ (slope). Diminishing antennal length had modest to no effects on wasp behaviour (Extended Data Fig. 2g).

We tested whether elongated antennae drive defensive behaviour in flies by pairing flies with wasps whose antennae had been removed. Strikingly, removing the wasp’s antennae abolished fly defensive responses, reducing them to a level comparable to that observed in pairings with control heterospecific flies (mean escape speed 0.11 and 0.03 mm/s, respectively; **Fig. 2c**, **Extended Data Fig. 2d,g**; **Supplementary Video 9**). Next, we paired flies with non-threatening heterospecific flies to which we added an elongated antenna-like filament. Heterospecifics with a synthetic antenna induced a modest but significant increase in defensive behaviour in *D. melanogaster* females compared to unmodified controls (**Fig. 2d**, **Extended Data Fig. 2e**; **Supplementary Video 10**). Lastly, we examined the influence of antennal length on defensive response by cutting the antennae of male wasps from a normal length of 13 segments to nine, six, three, or zero remaining segments. We found that fly escape speed and frontal-bias index scaled linearly with antennal length (**Fig. 2e**, **Extended Data Fig. 2f, g**). This relationship may explain our prior observation that male wasps, which have longer antennae than female wasps, elicit stronger escape responses in flies (**Fig. 1c, d, f**). Thus, elongated antennae are a conspicuous visual cue that drives innate defensive behaviour in flies.

### LC neurons mediate wasp evasion

We performed genetic silencing experiments to identify neuronal cell types involved in the visual detection of wasps and the orchestration of defence. We focused on the lobula columnar (LC) neurons, a family of approximately 50 cell types that function as visual feature detectors (**Fig. 3a**). Using available cell-type-specific split-GAL4 driver lines ^9,27,28^, we individually silenced 27 LC types by expression of tetanus toxin (TNT) to block synaptic release. We paired LC-silenced flies with male wasps and quantified two defensive metrics defined above, escape speed and frontal-bias index (**Fig. 3b**, **Extended Data Fig. 3a**). As a control for genetic background effects, we also included an empty split-GAL4 line with no expression in the nervous system in our screen ^29^. We computed the joint distance from the genetic control across the two defensive metrics for each LC-silenced genotype (see Methods). This analysis identified five LC types — LC9, LC15, LC21, LC25 and LPLC1 — whose silencing produced significant deficits in defence (**Extended Data Fig. 3b–d**). Thus, a small set of LC types are required for wasp-evoked defensive responses.

**Figure 3.**
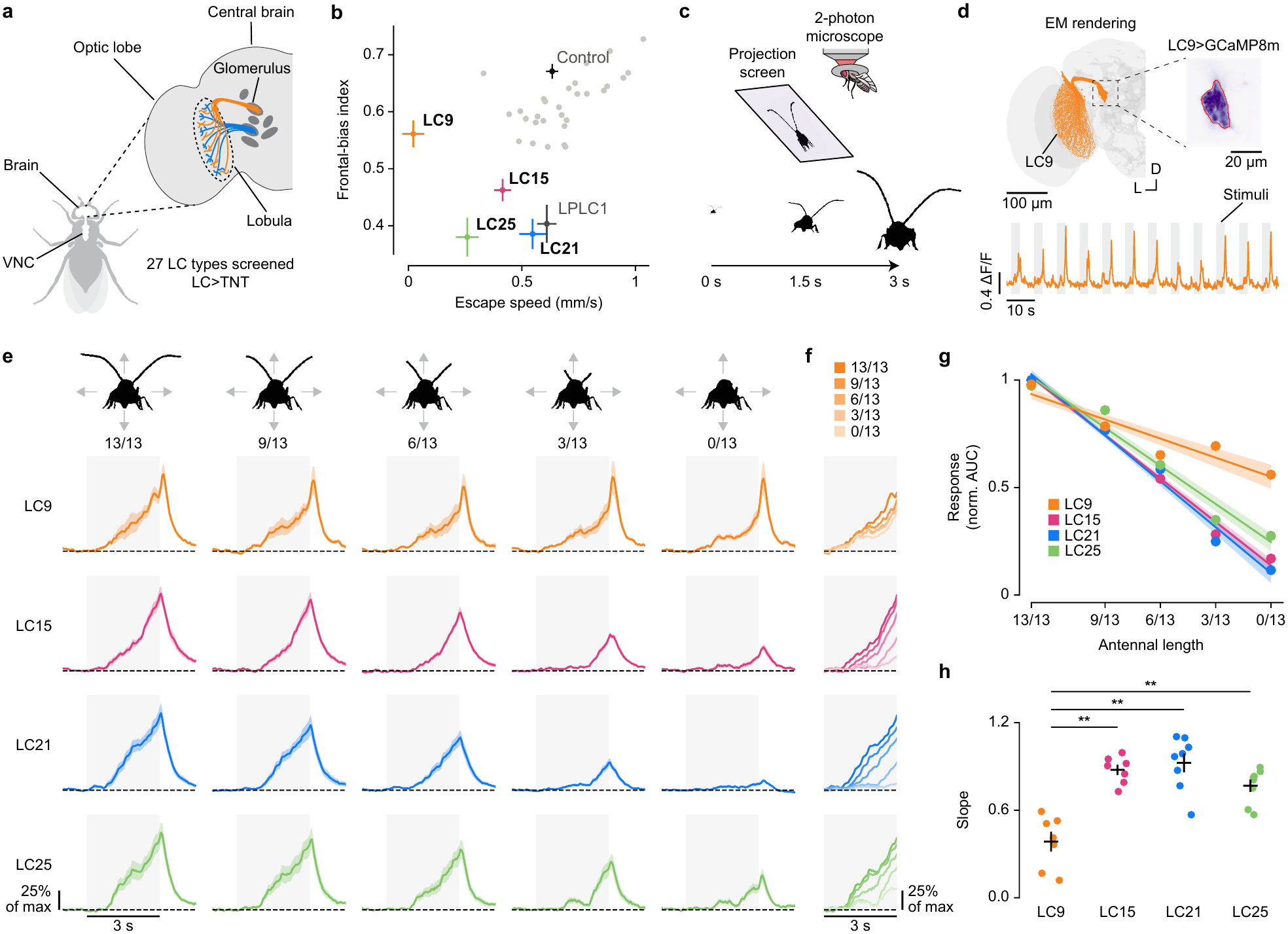
LC types required for wasp defence encode wasp antennal length. **a,** Each LC type has dendrites that tile retinotopic space in the lobula, and axons that converge to an optic glomerulus in the central brain. Tetanus toxin (TNT) was expressed in 27 individual LC types using cell-type-specific split-GAL4 lines as part of the silencing screen. VNC, ventral nerve cord. **b,** Escape speed versus frontal-bias index for each silenced LC type and the empty split-GAL4 control. The five LC types whose silencing significantly impaired defence (Extended Data Fig. 3b) are labelled. Other LC types are shown as grey dots. Crosses, mean ± s.e.m. along each axis. *n* (pairs; split-GAL4>TNT): LC9, 82; LC25, 57; LC15, 97; LC25 #2, 79; LC21, 58; LC11 #2, 81; LPLC2 #2, 50; LC11, 53; LC21 #2, 59; LC31b, 77; LC14, 52; LC33, 54; LC10a, 58; LPLC2, 65; LC6, 106; Control, 145; LC4, 44; LC10a-d, 89; LPLC1, 63; LC27, 69; LC4 #2, 42; LC15 #2, 23; LC18, 62; LC24, 65; LC38, 57; LC43, 82; LPLC4, 110; LC13, 60; LC22, 50; LC17, 45; LC26, 63; LC12, 54; LC16 #2, 50; LC16, 57; LC20, 65. All five LC types labelled here showed significant deficits for both defensive metrics in pairwise comparisons with their matched GAL4-only genetic control (Extended Data Fig. 3c, d). **c,** Individual flies were presented with movies of the silhouette of walking wasps while the activity of LC neurons expressing a calcium indicator was imaged with a 2-photon microscope. **d,** Top left: EM reconstruction of the LC9 population in the right hemisphere of the brain (orange). Top right: view of the LC9 glomerulus during an imaging session. Bottom: example ΔF/F trace from an LC9 glomerulus during stimulus presentation (grey bars). D, dorsal; L, lateral. **e,** Mean calcium responses of LC9, LC15, LC21, and LC25 (rows) to movies of walking wasp silhouettes expanding to simulate approach whose antennae were truncated to 13/13 (intact), 9/13, 6/13, 3/13, or 0/13 of their full length (columns). Representative frame of each stimulus variant shown at top. For each cell type, responses are normalized to the peak response to the intact (13/13) stimulus. Grey shading, stimulus period (0–3 s); dashed line ΔF/F = 0; line and shading, mean ± s.e.m. Here and in f–h, *n* (flies): LC9, 7; LC15, 7; LC21, 8; LC25, 8. **f,** Mean responses during the stimulus period (0–3 s) from e overlaid for each cell type. Shading indicates antennal length, from 13/13 segments (dark) to 0/13 segments (light). **g,** Integrated responses (area under the curve during the stimulus period, 0–3 s) as a function of antennal length. For each fly, responses were normalized to the maximum across the five antennal lengths. Points, mean across flies; lines and shading, mean ± s.e.m. of linear fits to individual flies. Same flies as in e. **h,** Slopes of the per-fly linear fits in g. Points, individual flies; crosses, mean ± s.e.m. Kruskal–Wallis test, *H* = 17.8, *P* = 4.76 × 10^−4^; two-sided Mann–Whitney *U* tests with Bonferroni correction: LC9 vs. LC15, *P* = 3.50 × 10^−3^; LC9 vs. LC21, *P* = 3.73 × 10^−3^; LC9 vs. LC25, *P* = 3.73 × 10^−3^; LC15 vs. LC21, *P* = 1.00; LC15 vs. LC25, *P* = 0.563; LC21 vs. LC25, *P* = 0.390.

### LC activity encodes wasp antennal length

Four of the cell types identified in our silencing screen have previously been shown to be tuned to small visual features: LC9, LC15, LC21, and LC25^10,11^. Activity in these four LC types may therefore encode visual features of the wasp. We tested this by expressing GCaMP8m in each of these LC types individually and recording calcium signals from their respective axonal glomeruli while head-fixed flies viewed movies of the silhouette of a wasp walking towards them (**Fig. 3c, d**; **Supplementary Video 11**). In the movies, we expanded the moving silhouette at a constant rate to simulate approach (18°to 146°of visual field). We found that all four LC types responded strongly to these naturalistic wasp stimuli (**Fig. 3e**, first column). Thus, the same neurons whose silencing impairs the fly’s defence are activated by the form of a wasp.

We determined whether the magnitude of the LC responses scaled with antennal length, potentially matching our observation that fly defensive behaviour scaled with antennal length (**Fig. 2e**). We generated movie variants in which the antennae of the wasp were shortened to 9/13, 6/13, 3/13, and 0/13 of their full length, paralleling the antennal manipulations in our behavioural experiments (**Fig. 3e; Supplementary Video 11**). The integrated response of all four cell types to these stimuli scaled linearly with antennal length, with longer antennae producing stronger activity (**Fig. 3f, g**). This relationship was preserved with non-expanding wasp movies and therefore does not depend on the visual properties of looming (**Extended Data Fig. 4a–d**; **Supplementary Video 12**). The parallel scaling of neural and behavioural responses with antennal length suggests that activity in these neurons regulates the intensity of defence.

### LC9 forms a composite representation of the wasp

The responses of LC15, LC21, and LC25 to the wasp stimulus movies were nearly eliminated when the antennae were removed, whereas the responses of LC9 were approximately halved (**Fig. 3g, h**). This suggests that LC9 may respond to other components of the wasp in addition to the antennae. We investigated this by developing a second panel of movies in which the wasp was deconstructed and presented as various subsets of its components, including the antennae, body, or both (**Fig. 4a**, **Supplementary Video 13**). LC15 and LC21 responded comparably whether the antennae were presented alone or together with the body, and LC25 responded only modestly less to the antennae alone. This further supports that the responses of these three LC types to wasps are driven predominantly by the antennae.

**Figure 4.**
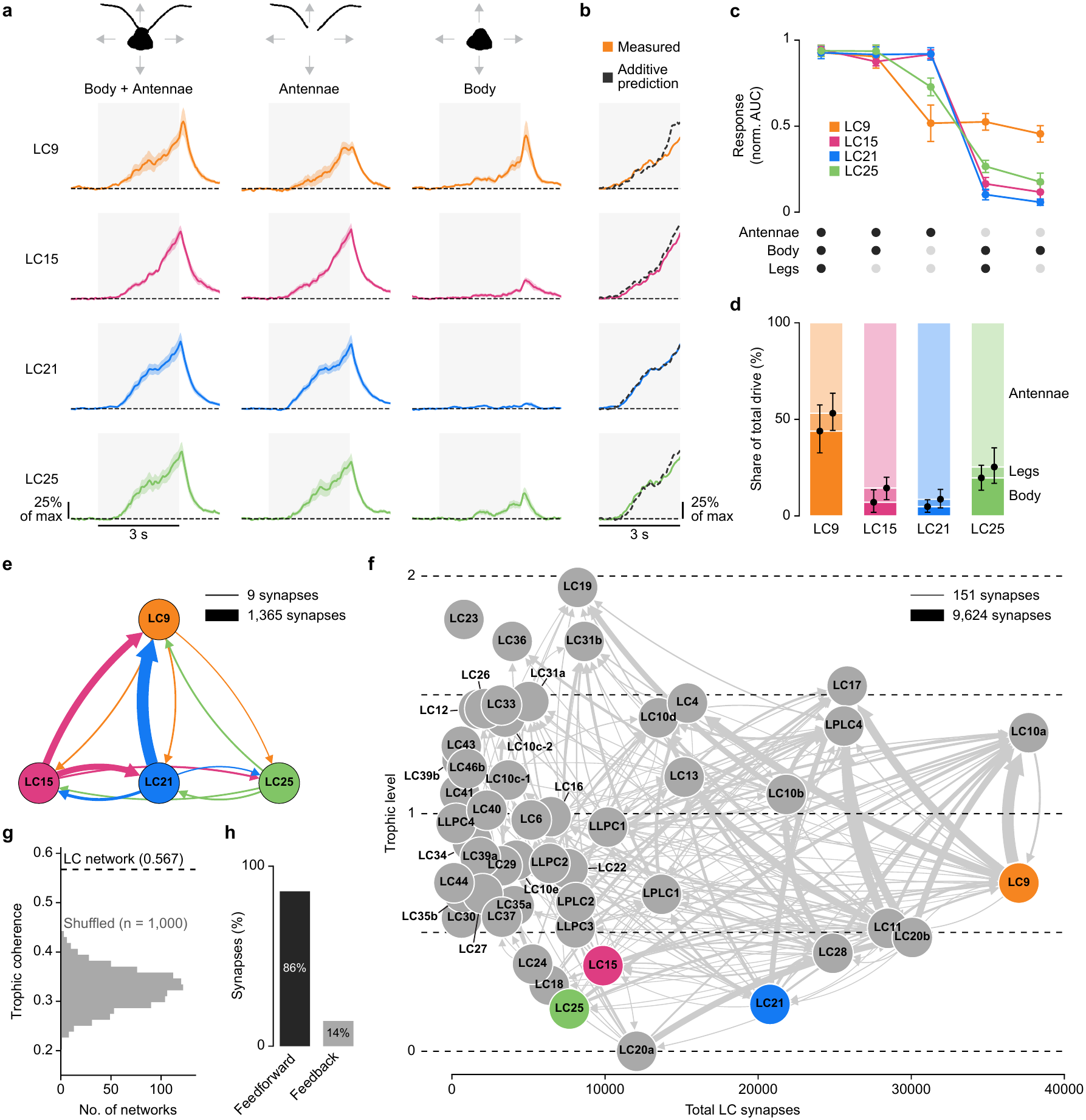
LC9 forms a composite representation of the wasp. **a,** Mean calcium responses of LC9, LC15, LC21, and LC25 (rows) to movies of approaching wasps showing the body and antennae together (left), the antennae alone (middle), or the body alone (right). Representative frame of the three stimulus variants shown at top. For each cell type, responses are normalized to the peak response to body + antennae stimulus. Grey shading, stimulus period (0–3 s); dashed line ΔF/F = 0; line and shading, mean ± s.e.m. Here and in b–d, *n* (flies): LC9, 7; LC15, 7; LC21, 8; LC25, 8. Data are from the same recordings as in Figure 3e–h. **b,** Measured response to the body + antennae stimulus (coloured; from left column of a) compared with the additive prediction (dashed black), computed for each fly as the sum of its responses to the antennae-alone and body-alone stimuli. LC9 measured versus additive prediction, ratio 0.97 (95% bootstrap CI 0.79–1.16). **c,** Integrated responses (area under the curve during the stimulus period, 0–3 s) to five stimuli containing different combinations of wasp components (dot matrix below; black, component present). The body + legs + antennae and body + legs stimuli are the intact (13/13) and 0/13 stimuli of Fig. 3e; the body + antennae, antennae and body stimuli are those shown in a. For each fly, responses were normalized to the maximum across the five stimuli. Points and error bars, mean ± s.e.m. across flies. Two-sided paired t-tests on per-fly integrated responses, Holm-corrected: antennae alone vs. body + antennae: LC9, *P* = 0.0214; LC15, *P* = 0.399; LC21, *P* = 0.471; LC25, *P* = 1.95 × 10^−3^; body alone vs. body + antennae: LC9, *P* = 0.0214; LC15, *P* = 2.15 × 10^−5^; LC21, *P* = 6.87 × 10^−4^; LC25, *P* = 1.95 × 10^−3^. **d,** Contribution of each wasp component to the integrated response of each LC type. For each fly, integrated responses to the five stimuli in c were fit with a non-negative additive model, and the fitted weights were expressed as shares of the total (see Methods). Stacked bars, mean share attributed to the body (dark), legs (intermediate), and antennae (light). Points and error bars, mean and 95% confidence interval of boundaries. Median *R*^2^ of the additive fit across flies: LC9, 0.846; LC15, 0.970; LC21, 0.991; LC25, 0.972. **e,** Synaptic connectivity between LC9, LC15, LC21, and LC25 (male CNS, lobula). Arrow width is proportional to synapse count. **f,** Trophic analysis of the LC-to-LC network. Each node is an LC type, positioned by its trophic level (y axis) and its total LC-to-LC synapses (input + output; x axis). Edge width is proportional to synapse count. For visual clarity, only connections of strength >150 synapses are shown. **g,** Trophic coherence of the LC-to-LC network (dashed line) compared with shuffled, degree-matched networks (grey histogram; 0.325 ± 0.038, mean ± s.d.). *z* = 6.3; empirical *P* < 0.001 (two-sided). **h,** Percentage of LC-to-LC synapses that are directed up from lower to higher trophic levels (feedforward; 86%), or down from higher to lower trophic levels (feedback; 14%).

LC9, however, responded maximally when the antennae and body components were presented together, and responded more weakly to the antennae or body alone (**Fig. 4a**). Summation of the responses to the individual component stimuli recapitulates the response to the two-component stimulus, both for the integrated response (**Fig. 4c**) and the instantaneous response (**Fig. 4b**). We quantified the contribution of each wasp component to the response magnitude of each cell type using linear regression. Here, we fit the integrated responses to a full panel of deconstructed stimuli, which included the wasp’s legs as a component. In LC15, LC21, and LC25, the antennae dominated, accounting for 75–91% of the response; whereas in LC9, the antennae and body contributed equally, accounting for 47% and 44% respectively (**Fig. 4d**). The legs contributed minimally across all four LC types. These relationships were preserved in non-expanding wasp movies (**Extended Data Fig. 5a–d**, **Supplementary Video 14**). Therefore, LC15, LC21, and LC25 represent a single component of the wasp, the antennae, while LC9 represents the sum of two components, the antennae and the body.

### LC-to-LC connections support multi-component object representations

What circuit mechanisms impart LC9 with a representation of two visually distinct components? LC neurons have been conceptualized as a single layer of feature detectors that derive their selectivity through inputs from intrinsic optic lobe neurons. An alternative possibility is that LC neurons receive their selectivity in part from connections with other visual projection neurons, including other LCs. LC9, for example, could inherit its antennal sensitivity from the more narrowly tuned antennal detectors: LC15, LC21, and/or LC25. We analysed connectivity among these four cell types in multiple connectomes and found recurrent, monosynaptic connectivity, with strong input from LC15 and LC21 onto LC9 (**Fig. 4e**; **Extended Data Fig. 6a**) and 15-fold weaker reciprocal connections from LC9. Both LC15 and LC21 are cholinergic ^11^ and therefore excitatory. This connectivity is consistent with a feedforward, hierarchical model in which LC15 and LC21 provide antennal-feature information to LC9. LC9 may combine this antennal information with other inputs carrying body-feature information to reconstruct a more complete representation.

LC-to-LC connections, arranged as a feedforward hierarchy, may be a more prevalent motif than previously appreciated. We examined all LC-to-LC connections and found rich interconnectivity among LC types, with each LC type receiving input from at least one other LC type (median = 6.7% lobular input from other LC types). We tested to what degree the entire LC-to-LC network could be characterized as having a feedforward, hierarchical organization by applying a trophic-coherence analysis, borrowed from the study of food webs. This analysis measures how consistently connections run from lower to higher levels; each node is assigned a position within the hierarchy such that nodes that primarily output onto others occupy lower levels, while nodes that primarily receive input occupy higher levels (see Methods). This analysis revealed that LC types occupy a distribution of trophic levels (**Fig. 4f; Extended Data Fig. 6b, c**). The LC-to-LC network was substantially more directed than shuffled, degree-matched networks (**Fig. 4g; Extended Data Fig. 6d**) and 86% of synapses pointed upward along the trophic axis (**Fig. 4h; Extended Data Fig. 6e**), indicating strong feedforward organization. Notably, the more narrowly tuned LC types we characterized — LC15, LC21, and LC25 — were assigned low trophic levels, and LC9 occupied a higher position. This hierarchical organization may govern the combination of simpler component-selective signals into more complex representations.

### Spatial filtering drives wasp-directed turning

The fly connectome provides the opportunity to ask how the visual representation of a wasp is transformed into defensive action. We searched across connectomes ^30,31^ for direct connectivity from the identified LCs onto premotor descending neurons (DNs) ^32^, which connect the brain to motor circuits in the ventral nerve cord (**Fig. 5a**). The strongest of any direct connections was from LC9 to DNp09 (**Fig. 5b; Extended Data Fig. 7a**). The LCs selective for the antennae (LC15, LC21, and LC25) also connect indirectly to DNp09 via LC9 and two additional LC types that connect directly with DNp09, LC31a and LC31b (**Fig. 5c, d; Extended Data Fig. 7b–d**). Together, LC9, LC31a, and LC31b account for nearly 20% of all DNp09 input synapses (**Fig. 5e; Extended Data Fig. 7e**), indicating that these three cell types substantially shape DNp09 activity.

**Figure 5.**
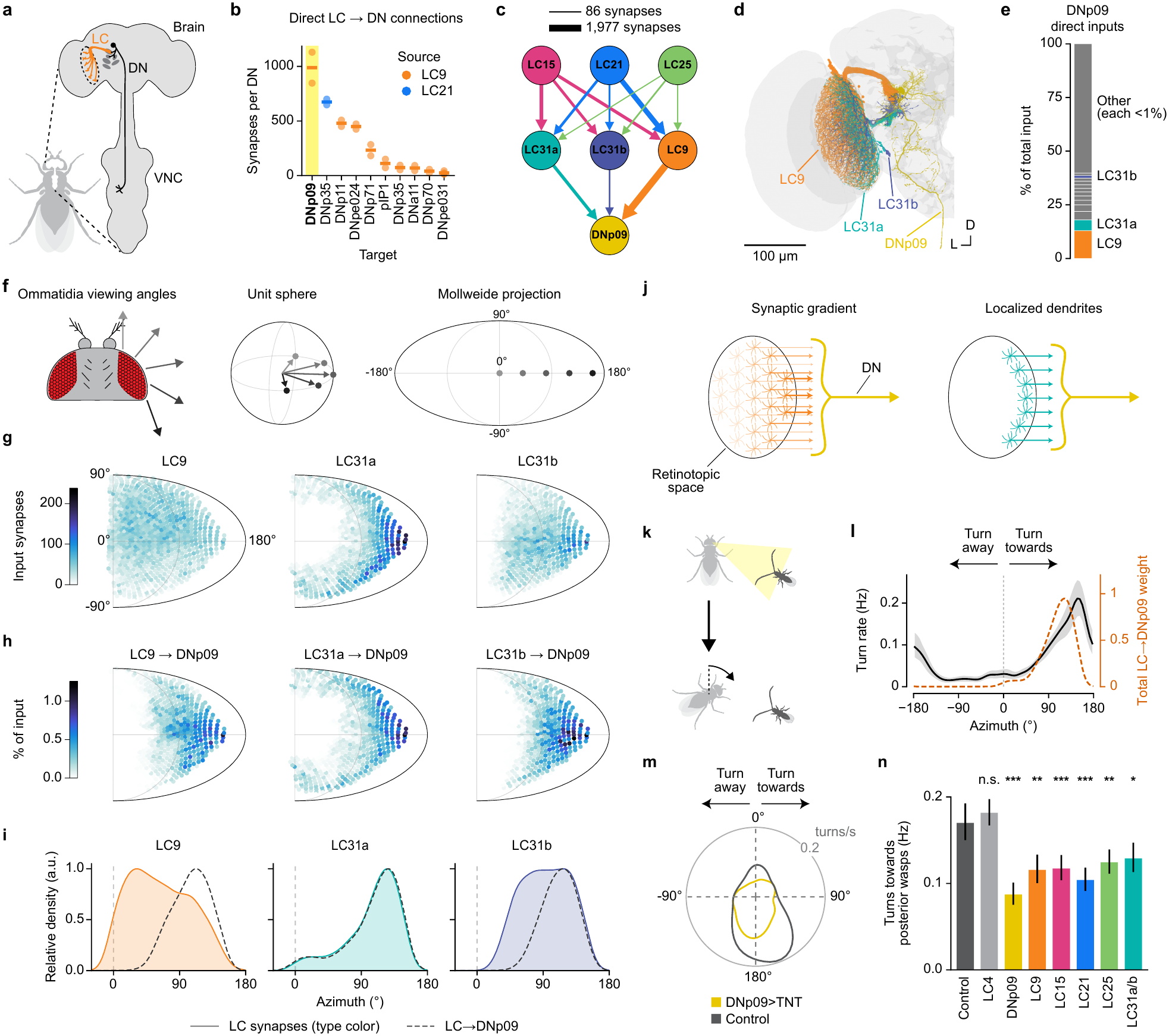
Spatial filtering of LC input to DNp09 directs turning towards wasps. **a,** Premotor descending neurons (black) relay information from the brain to the ventral nerve cord (VNC). Some descending neurons synapse directly with LC neurons (orange) at their optic glomeruli. **b,** Input synapses per descending neuron downstream of the imaged LC types. Colour indicates the source cell type (LC9, orange; LC21, blue). Points, individual descending neurons; lines, mean. DNp09 has the highest average input strength and is highlighted. **c,** Connectivity from antennal-feature encoding LC types (LC15, LC21, and LC25) onto DNp09 through LC9, LC31a, and LC31b. Edge width is proportional to synapse count. **d,** EM reconstruction of all LC9, LC31a, LC31b, and DNp09 neurons in the left hemisphere. D, dorsal; L, lateral. **e,** DNp09 inputs ranked by the proportion of contributed input synapses (15,235 synapses across both DNp09 neurons). LC9, 13.0%; LC31a, 5.0%; LC31b, 1.2%. Any inputs that individually contribute <1% of synapses are pooled as Other. **f,** Illustration of the relationship between ommatidial viewing angles on the eye of the fly (left), vectors on a unit sphere (middle), and points on a Mollweide projection (right). **g,** Predicted receptive fields of LC9, LC31a, and LC31b populations, shown as Mollweide projections. Each point is an ommatidium positioned at its viewing angle and coloured by the number of input synapses from the corresponding lobula column. The map for the left eye was mirrored about the 0°–180°azimuthal meridian and averaged with the right eye. **h,** Receptive fields inherited by DNp09 from LC9, LC31a, and LC31b, shown as Mollweide projections. Each point is an ommatidium positioned at its viewing angle and coloured by the percentage of input contributed by the corresponding lobula column, with each LC type’s contributions across space summing to 100%. The map for the left eye was mirrored about the 0°–180° azimuthal meridian and averaged with the right eye map. **i,** Azimuthal distribution of the predicted receptive field for each LC type (solid, coloured), and the contribution of that type to the receptive field of DNp09 (dashed). Each distribution is normalized to its peak. **j,** Two mechanisms for producing spatially directed behaviours. Left, synaptic gradient: a visual projection neuron population (orange) with dendrites that tile retinotopic space carries information from the whole visual field, but a gradient in output synapses across cells (arrow thickness/opacity) results in a spatially restricted receptive field in a downstream descending neuron (yellow). Right, localized dendrites: a visual projection neuron population (teal) with spatially restricted dendritic arbors carries information from a restricted region of space; a synaptic gradient is therefore not needed. **k,** Schematic of the wasp-directed turning behaviour orchestrated by the LC–DNp09 microcircuit described in this figure: a fly turns towards a wasp that is in its lateral-posterior visual field (yellow). **l,** Turning rate as a function of wasp’s azimuthal position relative to the fly (black) for wild-type flies paired with male wasps in a circular chamber, overlaid with the combined LC9, LC31a, and LC31b input to DNp09 (orange dashed; normalized to peak). Left turns were mirrored about the 0°azimuthal axis such that positive azimuths indicate turns towards the wasp. Shading, 95% bootstrapped confidence interval. *n* = 1,574 turns from 57 flies. Binomial GLM of turn counts on the connectomic prediction: slope 1.94 (95% CI 1.65–2.22, cluster-robust over flies), *P* = 1.3 × 10^−19^; *r* = 0.75. **m,** Turning rate as a function of wasp’s relative azimuth position for flies expressing TNT in DNp09 (yellow) and empty split-GAL4 control (grey), paired with male wasps in a circular chamber. *n* (turns): Control, 1,583; DNp09>TNT, 1,216. **n,** Rate of turns towards wasps in the lateral-posterior visual field (121–177°azimuth) for empty split-GAL4 control flies and flies expressing TNT in the indicated cell types. Error bars, Wilson 95% confidence intervals. *χ*^2^ tests on pooled counts compared with the empty split-GAL4 control, Holm-corrected across all 28 pairwise comparisons: LC4, *P* = 1.00; DNp09, *P* = 1.24 × 10^−10^; LC9, *P* = 1.09 × 10^−3^; LC15, *P* = 6.76 × 10^−4^; LC21, *P* = 1.46 × 10^−6^; LC25, *P* = 4.10 × 10^−3^; LC31a/b, *P* = 0.0398. *n* (turns): Control, 245; LC4, 548; DNp09, 177; LC9, 191; LC15, 247; LC21, 227; LC25, 304; LC31a/b, 219.

Activation of the DNp09 neuron in one hemisphere drives ipsilateral turning ^33^. DNp09 may thus relay the motor commands that cause flies to turn towards wasps in the lateral-posterior visual field (as described in **Fig. 1k, l**). This spatially selective visuomotor transformation requires that DNp09 receive wasp-feature input preferentially from the lateral-posterior region of visual space. We tested this by mapping the columnar inputs of each LC type onto ommatidial viewing directions in visual space ^26^ (**Fig. 5f**; see Methods), which allowed us to predict the spatial receptive field of each LC neuron and cell type (**Fig. 5g**). Weighting each LC neuron’s receptive field by its synaptic weight onto DNp09 then gives the receptive field that DNp09 inherits from each LC type. This analysis revealed two distinct spatial filtering mechanisms. As a population, LC9 tiles all visual space (**Fig. 5g; Extended Data Fig. 7f**). DNp09, however, receives input predominantly from LC9 neurons with posterior receptive fields (**Fig. 5h, i**), consistent with a wiring motif termed synaptic gradients ^34^. LC31a and LC31b achieve the same spatial filtering onto DNp09 by a different logic. The dendrites of LC31a/b are confined to the posterior lobula (**Fig. 5g; Extended Data Fig. 7f, g**), and therefore the LC31a/b populations can only convey information from the lateral-posterior visual field to DNp09 (**Fig. 5h, i**). Thus, synaptic gradients and retinotopically localized dendrites are mechanisms that together limit DNp09’s wasp-feature input to the lateral-posterior visual field (**Fig. 5j; Extended Data Fig. 8a–c**).

This filtering mechanism may account for the spatial organization of wasp-directed turning behaviour (**Fig. 5k**). We predicted the receptive field of DNp09 from the combined inputs of LC9 and LC31a/b and compared it with the fly-to-wasp turning “behaviour field” — the distribution of wasp azimuths at which flies initiated turns (**Fig. 5l**). The connectomic prediction accounted for much of the spatial structure of wasp-directed turning (binomial GLM, *P* = 1.3 × 10^−19^; 81% of the performance of an unconstrained fit; **Extended Data Fig. 9a, b**). We tested whether genetically silencing individual cell types within this LC–DNp09 microcircuit would impair the fly’s ability to turn towards wasps positioned in its lateral-posterior visual field. We paired individual LC- or DN-silenced flies with a male wasp in a circular behavioural chamber. We found silencing any cell type within the microcircuit significantly reduced turning towards wasps in the lateral-posterior visual field (**Fig. 5m, n; Extended Data Fig. 9c, d**), with DNp09 silencing causing the largest magnitude impairment. In comparison, silencing LC4, an LC type not identified as part of the LC–DNp09 microcircuit, did not impair turning. These results define a circuit that builds a composite representation of a wasp, spatially filters and transmits that representation downstream to a premotor pathway, resulting in the fly turning towards a wasp that moves behind it.

## Discussion

Parasitoid wasps are pervasive natural enemies of *Drosophila* and elicit diverse behaviours that are thought to reduce the risk or consequences of parasitism ^35^. Although multiple modalities contribute to wasp detection in flies, vision is a major driver of sustained behavioural changes, most notably in egg-laying and courtship ^3–5,36^. We observed that flies also respond acutely to wasp encounters with a dynamic suite of visually guided defensive actions that allow the fly to evade physical contact. This reaction suggests that the wasp is considered an immediate threat to the fly itself, which is unexpected given that the female wasps used in our experiments attack pupae or larvae, while the male wasps are non-predatory. The expression of this defensive repertoire in flies is triggered by the sight of the wasp’s elongated antennae, and the intensity of defence scales with antennal length. This aligns with the classic ethological concept of a sign stimulus, in which a specific sensory feature of an animal or object in the environment triggers an appropriate innate behavioural response, often in proportion to the magnitude of that feature ^37^. Elongated antennae are also characteristic of other sympatric predators of flies that do pose an immediate threat, such as ants, and may therefore represent a heuristic cue for defence against a broader class of threatening animals.

The threatening association of elongated antennae could impose selection in the predators for morphologies or behaviours that minimize their detection by prey. For example, female parasitoid wasps of several species, including all three studied here, exhibit reduced antennal length relative to non-predatory conspecific males. *Syntretus perlmani*, a parasitoid of adult *Drosophila*, appears in published videos ^38^ to reposition its antennae backward during close pursuit and attack, potentially obscuring its antennae from view. Conversely, non-threatening species may exploit the same cue association to appear threatening and gain protection through Batesian mimicry. For example, stilt-legged flies and some species of jumping spiders extend their forelegs forward in a conspicuous manner that resembles the elongated antennae of wasps and ants ^39,40^. Together, these examples suggest that elongated antennae may constitute a broadly recognized signal of threat subject to opposing selective pressures across taxa.

In the brain, we found four LC visual projection neuron types that respond to the sight of an approaching wasp. These cell types are also recruited in other visual contexts. Three of the four LC types identified in the present study — LC15, LC21, and LC25 — respond to thin moving bars ^10,11^, and LC15 is implicated in distance estimation during gap crossing ^41^. The fourth, LC9, is thought to be tuned to small moving objects and has been linked to the engagement of conspecifics during courtship and aggression ^8,33,42^. Conversely, wasps are likely to recruit additional LC types beyond these four. Indeed, LC4, an LC type whose silencing did not alter the defensive behaviours examined here, has been suggested to mediate wasp-induced changes in reproductive behaviour ^3,5^. Collectively, these observations are consistent with the idea that visual objects are represented in a population code distributed across LC types, with LCs recruited in different combinations for different behaviours ^8,11^. Nonetheless, our silencing screen indicates that it is a small subset of LCs that convey the information essential for acute defence against wasps.

How stimulus selectivity arises is unknown for most LC types (for an exception, see ^29^). For LC15, connectome analysis predicts that direct input from the medulla may give rise to orientation-invariant selectivity for thin bars ^43^. Our data suggest that more complex tuning in some LCs can be assembled by combining input from intrinsic neurons of the optic lobe with input from other LCs. Unlike the LCs that selectively represent the antennae, LC9 represents both the wasp’s antennae and body. Direct input from LC15 and LC21 suggests that LC9 inherits its antennal sensitivity from these more selective cell types. Our analysis of the broader LC network suggests that feedforward LC-to-LC connectivity may be essential for the generation of richer and perhaps more ethologically meaningful representations by progressively combining the simpler feature selectivity of certain LC types to construct more complex tuning profiles in others.

The logic of object representation described here is reminiscent of face coding in the primate brain. Face-selective neurons in the macaque inferotemporal cortex are tuned to constituent facial features, such as inter-eye distance, iris size, or aspect ratio ^22^. In face cells, as we observed with LC tuning for antennal length, responses scale linearly with the magnitude of the feature ^22,24^. In the primate, it is hypothesized that more specific, feature-selective channels feed onto combinatorial channels: posterior face patches respond to isolated features ^44^, and responses become progressively more holistic along the visual hierarchy ^23^. Furthermore, face cells tuned to multiple features combine constituent responses near-linearly ^22,44^. This combinatorial coding mirrors the summation of antennal and body representations performed by LC9, and we present a synaptically resolved implementation of this longstanding hypothesis underlying visual object reconstruction from constituent features.

The appropriate response to an object depends not only on its identity but also its location. Consistent with this, flies exhibited a repertoire of directed defensive actions whose expression varied with the position of the wasp within the fly’s visual field. For example, flies orient towards a wasp that threatens to move out of sight behind the fly. Our connectomics analysis reveals how visual information about wasp identity and location is translated into this act of vigilance. We find that, although LC9 tiles visual space, DNp09 does not sample the LC9 population uniformly. Instead, DNp09 receives input preferentially from LC9 neurons with posterior receptive fields. This spatial filtering of LC9’s wasp representation results in turning when the wasp is in a more threatening posterior location. The organization of this visuomotor pathway resembles that of the vertebrate tectum and superior colliculus. In these brain structures, retinotopically organized inputs tuned to diverse visual features are coupled to motor outputs according to the properties and location of a stimulus ^14,45–49^. The fly connectome reveals the precise logic of such visuomotor transformations at synaptic resolution; the spatial filtering mechanisms we identified for turning may extend to the other defensive behaviours we observed. Each behaviour may be produced by a different spatial filter such that wasp detection in each spatial location is coupled to the best-suited action.

Together, these findings reveal a circuit logic for visually guided defence against a natural enemy: visual features associated with distinct morphological components are extracted and recombined to build a behaviourally meaningful representation of the predator, which is then filtered according to its location to drive context-appropriate defensive action.

## Methods

### Insect stocks and husbandry

Flies and parasitoid wasps were reared at 25°C and 55% relative humidity on a 12-h light/dark cycle. Wasp cultures were maintained as described elsewhere ^50^. Briefly, vials containing cornmeal-agarose food were populated with 20–30 adult flies for 2–3 days to allow for egg-laying. After removal of the adult flies, 7–10 mated female wasps were introduced to parasitize the developing fly offspring, with 3 ml 50% honey-water provided on the cotton plugs as a food source for the wasps. For detailed animal sources and genotypes, see Supplementary Table 1.

### Behavioural experiments

#### Pairwise interaction assay

All behavioural experiments were performed using 3–7-day-old flies and parasitoid wasps. Experimental fly crosses were maintained in vials containing molasses-agarose food. At least two days before experiments, flies were genotyped under CO_2_ anaesthesia and transferred to a vial containing cornmeal-agarose food. At least two days before experiments, wasps were sexed under CO_2_ anaesthesia and transferred to vials containing cornmeal-agarose food, supplemented with 3 ml 50% honey-water on the cotton plug. Female Canton-S *D. melanogaster* were used as wild-type flies in behavioural experiments. Male *D. micromelanica* were used as heterospecific flies in behavioural experiments. The UAS-TNT-E line used in genetic silencing experiments was backcrossed to Canton-S for six generations. We performed experiments with two distinct split-GAL4 driver lines for a subset of LC types (7 of 27 types) in genetic silencing experiments.

For quantifying behaviour in high resolution, multiple animal pairs were filmed in parallel within custom 3D-printed assemblies secured between two pieces of 0.25 mm transparent acrylic. For experiments performed in rectangular chambers, the assembly consisted of 30 chambers each with two connected compartments: a large compartment (5.0 mm tall × 19.6 mm wide) for observing interactions and a small compartment (5.0 mm tall × 5.0 mm wide) accessible via a 0.6 mm gap that only wasps could pass through, allowing for intermittent periods without interaction. For experiments performed in circular chambers, the assembly consisted of 16 chambers (13.0 mm diameter). Chamber depth varied depending on the partner animal: 1.8 mm for female wasps or heterospecific flies, or 2.6 mm for male wasps. These differences in depth were to accommodate variation in the vertical profiles of the partner animals, including antennal length, and to ensure the two animals could not pass above one another without making contact.

Animals were anaesthetised on ice and aspirated into individual chambers of a chilled temporary assembly. Once fully loaded, the animals were transferred into a room-temperature experimental assembly and secured with an acrylic lid. The assembly was then moved to a light-tight video-recording setup in a temperature- and humidity-controlled room. Within the setup, the assembly was placed atop an infrared panel light (850 nm; Smart Vision Lights), centred beneath a white low-angle square light (6000 K; FPQ3-120SW, CCS), and video recordings were performed using a monochrome USB3 camera (Basler acA2440-75um) attached to a 0.25× telecentric lens (Edmund Optics #55-349) at 40 Hz via pylon Viewer software (Basler). In darkness, the animals were given 10 min to recover from cold anaesthesia and habituate to the chamber. Subsequently, animals were filmed for 16 min under infrared illumination (“dark” condition), followed by 31 or 61 min filmed under white-light illumination (“light” condition).

#### Synthetic antenna experiments

Male heterospecific flies (*D. micromelanica*) were anaesthetised using CO_2_ and secured in place dorsal-side up on the CO_2_ diffusion pad. Filaments approximately 2.3 mm in length were cut from the tip of a paintbrush (Princeton Neptune Series 4750 Round 1). A single filament was secured to the tip of a vice, the exposed end dipped into a small droplet of UV-curable glue (JDiction low viscosity UV resin hard type), and then secured in a clamp affixed to a micromanipulator. The filament was positioned such that the exposed end gently made contact with the fly’s head, directly in between its antennae, and was secured by applying UV light (LED-200, Electro-Lite) for 60 s. The fly was allowed to recover for at least 2 days prior to being used for experiments. Both control and modified heterospecific flies were starved for 24 hours prior to being used for experiments to increase their activity.

#### Behavioural data analysis

All behavioural analysis was performed using custom-written code in MATLAB (Mathworks). DeepLabCut ^25^, a key-point detection algorithm, was used to track four points on flies (the ocellus and antennae-base midpoint on the head, a central point at the juncture of the thorax and abdomen, and the abdomen posterior tip) and five points on wasps (the tip of each antenna, the antennae-base midpoint on the head, a central point at the juncture of the thorax and abdomen, and the posterior tip of the abdomen). The central midpoint was used to track the animals’ x-y position, and their velocity was estimated as the distance travelled in x-y coordinates across 8 frames (200 ms). For determining the inter-animal distance, video frames were binarized to isolate discrete fly and wasp silhouettes. The inter-animal distance was determined as the shortest distance between silhouettes, and contact was defined as frames where the two silhouettes overlapped.

In the rectangular chamber assay, a given pairing was included for subsequent analysis if the partner animal was in the main compartment (“time in chamber”) for at least 5 min and crossed the chamber midline at least 20 times. Only frames in which the partner animal was in the main chamber were considered for analysis. For the circular chamber assay, pairings were included if the partner animal’s average velocity was at least 0.6 mm/s over the entire session. All inclusion thresholds were halved for the dark session.

Forward-running, backing-up, crabwalking, jumping, and turning events were identified based on empirically derived manual thresholds. Forward-running, backing-up, and crabwalking events were defined as sustained periods of directional movement that exceeded a baseline velocity threshold for at least 400 ms and reached a corresponding peak velocity within the epoch. Forward runs required a forward velocity greater than 0.075 mm/s with a peak greater than 0.75 mm/s. Conversely, backing-up events required a forward velocity less than −0.075 mm/s with a peak less than −0.75 mm/s. Crabwalks were defined similarly using lateral velocity, requiring a lateral velocity exceeding ±0.075 mm/s and a peak lateral velocity greater than ±0.75 mm/s in the same direction. Jump onsets were identified when 3-frame-smoothed forward acceleration crossed a threshold of 11.4 mm/s^2^ and forward velocity exceeded 1.245 mm/s within 125 ms. Jump offset was defined as the first subsequent point at which 10-frame-smoothed forward velocity fell below 0.825 mm/s. Jumps lasted at least 150 ms and met the following two criteria within the epoch: (1) reached either an angular speed greater than 225°/s or a forward velocity greater than 1.425 mm/s, and (2) reached either a peak forward acceleration greater than 21 mm/s^2^ or a peak angular acceleration greater than 7000°/s^2^. Turns were identified while flies were on the chamber floor and excluded forward-running, backing-up, and jumping epochs. Frames with forward velocity greater than 0.5 mm/s were additionally excluded. On the remaining frames, turns were defined as epochs in which the 10-frame-smoothed angular velocity exceeded 60°/s in either direction for at least 200 ms. Forward-running, backing-up, and turning events whose onset was within 5 s following a jumping epoch were excluded. Actions were considered contact-triggered if their onset occurred during or within 50 ms following frames where both animals’ silhouettes overlapped after binary thresholding.

A combination of inter-part angles and body pixel intensities were used to determine which surface of the chamber the fly was on — the sidewall, ceiling, or floor — and to identify transitions between surfaces, which were excluded from this classification. The fly was determined to be on the chamber sidewall if the magnitude of the angle formed between the line segment connecting the central midpoint to antennae-base midpoint and the line segment connecting the central midpoint to ocellus was greater than 11°. The fly was determined to be on the floor or ceiling if the magnitude of this angle was less than 6°. Frames where the fly’s long axis was not perpendicular to the floor (i.e. transitioning between surfaces) were excluded from wall and ceiling categories. These frames were identified when the length of the line segment connecting the antennae-base midpoint and the abdomen posterior tip was less than 90% of the peak of the distribution of line segment lengths (binned by 12.5 µm) observed for the entire session. Frames were then classified as either floor or ceiling categories in lights-on sessions depending on the mean pixel intensity of the segment connecting the central midpoint to ocellus, with floor frames having mean intensities greater than 72 (i.e. darker mean intensities correspond to floor frames). In the dark session, a DeepLabCut model trained to track the midpoint of the A3 abdominal stripe (only visible when the fly was on the floor) was used to distinguish whether the fly was on the floor or ceiling.

Fly escape speed was determined by the change in inter-animal distance between the fly at t = 0 ms and t = 200 ms and the partner animal’s location at t = 0 ms, for all frames satisfying the following criteria: partner animal velocity was 2–3 mm/s (or 1–3 mm/s in synthetic antenna experiments), inter-animal distance was less than 3 mm, and neither inter-animal contact nor a jump epoch had occurred for at least 5 s. The heading of each animal was determined by the line segment connecting the central midpoint and the antennae-base midpoint on the head. The central midpoint on the partner animal was used to determine the relative azimuthal position.

Frontal-bias index was calculated as the difference between the number of frames in which the partner animal was located in the frontal viewing field (azimuth 330–30°) and the number of frames in which the partner was located in the rear viewing field (azimuth 150–210°), normalized by the total number of frames in which the partner occupied either viewing field. Only frames meeting all of the following criteria were included: the fly was on the chamber floor or ceiling; fly velocity was less than 0.5 mm/s; partner animal’s velocity was greater than 0.5 mm/s (or greater than 0.125 mm/s in synthetic antenna experiments); neither inter-animal contact nor a jump epoch had occurred for at least 5 s.

To generate partner positional heatmaps, turn density heatmaps, and iso-contour maps, partner animal 2D spatial locations were binned into 0.2 mm square bins and smoothed using a two-dimensional Gaussian filter with *σ* = 0.9 mm. Partner positional heatmaps (Fig. 1h) were restricted to frames in which the fly’s velocity was less than 0.5 mm/s and the partner animal’s velocity was greater than 0.5 mm/s. Data were computed from frames and events pooled across flies of each pairing.

For turn-rate calculations, wasp-positional data from left turns were mirrored about the 0°–180° azimuthal axis, such that all turns toward the wasp (ipsiversive turns) corresponded to the right visual field. Turn rates were computed from turns and frames pooled across flies of each genotype. Circular turn-rate plots (Fig. 1l, Fig. 5m, Extended Data Fig. 9c) were smoothed with a circular Gaussian kernel (*σ* = 10°). The lower and upper bound of the lateral-posterior azimuthal range used to calculate turning rates (Fig. 1m, Fig. 5n) were set by the peak of the LC-to-DNp09 synaptic influence (121°; orange dashed trace in Fig. 5l) and the most posterior ommatidial position (177°), respectively.

Linear models (Fig. 2b, Extended Data Fig. 2c) were fit using measurements from all pairings, with predictor variables z-scored prior to fitting. Regression coefficients and associated *P* values were estimated from these models. Model performance was assessed by calculating *R*^2^ across group-mean behavioural values.

To identify lines of interest from the LC-silencing screen, escape speed and frontal-bias index were each z-scored using the mean and standard deviation pooled across all fly–wasp pairs of all split-GAL4>TNT groups, placing both metrics on a common scale. Each split-GAL4>TNT group was compared against the empty split-GAL4 control (SS01062>TNT) in this two-dimensional space. Effect sizes were computed as the Mahalanobis distance between the group’s centroid and the control centroid, 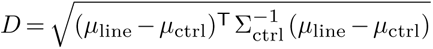 where Σ_ctrl_ is the covariance of the control flies. Significance was assessed with a two-sample permutation test on Hotelling’s *T* ^2^ statistic (pooled covariance). *P* values were corrected across all lines with the Holm–Bonferroni method. Defensive behaviour was considered altered relative to control for a split-GAL4>TNT group if its adjusted *P* value was below 0.05 and its *D ≥* 1.5. Silencing of an LC type was considered to alter defensive behaviour if at least one split-GAL4>TNT group specific to that LC type met these criteria.

### Functional imaging

#### Preparation and imaging

Female flies (4–10 days old) expressing GCaMP8m in the targeted LC type were anaesthetised on ice and mounted with UV-curable glue (Bondic) to custom stainless-steel fly-plates similar to those previously described ^51^. The proximal portion of the proboscis was glued to reduce movement. After recovery, the fly-plate was filled with saline (103 mM NaCl, 3 mM KCl, 5 mM TES, 8 mM trehalose, 10 mM glucose, 26 mM NaHCO_3_, 1 mM NaH_2_PO_4_, 1.5 mM CaCl_2_ and 4 mM MgCl_2_, adjusted to 270 mOsm, equilibrated to 7.3 pH). The cuticle of the posterior head capsule was dissected with a sapphire-tipped knife (World Precision Instruments) and forceps, and trachea over the posterior central brain were removed. Imaging was performed on an Ultima Investigator Plus 2-photon laser scanning microscope (Bruker) with a mode-locked, dispersion-compensated laser (Spectraphysics) tuned to 920 nm (15–30 mW power at the objective) and a 20×, 1.0 NA objective (Olympus). Emission was detected by a GaAsP photomultiplier tube (Hamamatsu) behind a 509/22 nm bandpass filter (Semrock). A single plane midway through the depth of the target LC glomerulus, ipsilateral to the stimulus, was imaged at 20–25 Hz.

#### Visual display

Stimuli were projected by a DMD projector (Wintech PRO4500) coupled by a liquid light guide (Thorlabs LLG5-4T) to a custom multi-source light beam (375–617 nm). The source was controlled by NI DAQ boards through a custom LED driver, allowing LEDs to be turned on during the return period of the x-scanning mirror in the 2-photon microscope ^51^. The projector was operated such that each of the 24 RGB bit planes was treated as a separate image to achieve a monochrome image stream. Movies were presented at 240 fps, with each frame displayed for 6 consecutive bit planes. Images were projected onto an acetal film screen (150 × 94 mm; 135.3°× 84.6°; McMaster 5742T51) positioned 63.5 mm from the fly at 45°azimuth on the side of the recorded hemisphere. Images were cylindrically warped relative to the fly’s viewing position. The background was a “fly-white” made from a mixture of LEDs whose relative intensities were set to excite the colour-sensitive photoreceptor types approximately equally ^52^. Wasps were rendered as black pixels against this white background.

#### Visual stimuli

Isolated male wasps (*L. heterotoma*) were filmed in a custom 3D-printed assembly of four chambers (6.8 mm tall × 6.8 mm wide × 12 mm deep), secured between two glass coverslips. The assembly was placed in front of a green panel light (530 nm; Smart Vision Lights) and filmed at 200 Hz using a USB3 camera (Basler acA2040-90umNIR), a 0.377× telecentric lens (Edmund Optics #34-015), and pylon Viewer software (Basler). From these recordings, three segments of 2.5–3 s were selected, each from a different wasp, in which the wasp traversed the chamber parallel to the camera axis. Segments of 2.5 s were extended to 3 s by prepending their first 0.5 s in reverse order. Videos were upsampled to 240 Hz for the projector. A binary threshold was applied to the raw videos to isolate the wasp silhouette from background pixels, and videos were then cropped by a common bounding box large enough to encompass all three video segments. This bounding box was then scaled and centred within an all-white canvas.

For each of the three source videos, we generated eight stimulus variants. These consisted of the intact wasp (13/13 antennal segments, or antennae + body + legs), three stimuli with shortened antennae (9/13, 6/13, and 3/13 antennal segments), and four stimuli with missing components (antennae + body, antennae, body, and body + legs or 0/13 antennal segments). Each variant was presented in two forms. In expanding stimuli, the silhouette’s bounding box was scaled linearly from 0.34× to 2.71× of its original size over the course of 3 s, corresponding to an expansion rate of approximately 42.6°/s along the diagonal, such that the stimuli expanded from approximately 18.1°to 145.8° of visual space along their diagonal. Across the expansion, the bounding box was positioned such that the centre of mass of the wasp’s body was anchored to a fixed point. In non-expanding stimuli, the bounding box was scaled by 2×, such that the stimuli were presented at approximately 107.5°along their diagonal. These stimuli were also positioned according to the centre of mass of the wasp’s body. Expanding and non-expanding stimuli were presented in separate, back-to-back recordings. Every fly was shown both stimulus sets. Each recording began with 10 s of a blank white screen, followed by three repeats of every variant from each source movie in randomized order (3 repeats × 3 source movies = 9 presentations of each stimulus type). Stimuli lasted 3 s, and successive stimuli were separated by 5 s of white background. Stimulus presentation was controlled with custom Python software.

#### Data analysis

Each recording was motion-corrected with piecewise-rigid registration in CaImAn ^53^. ROIs were automatically segmented from the mean frame of each recording using Otsu’s method. The fluorescence trace for each recording was the mean pixel value within the ROI for each frame. For each stimulus presentation, we computed ΔF/F = (F − F_0_)/F_0_, where F_0_ was the mean fluorescence during the 0.5 s before stimulus onset. Traces were linearly interpolated onto a common time base relative to stimulus onset at 20 Hz. For each fly, presentations of the same stimulus variant were averaged. Mean traces were then computed across flies. For display, traces were normalized to the peak of the mean response to a reference stimulus (Fig. 3, intact stimulus; Fig. 4, body + antennae stimulus). Where integrated responses are reported, they were defined as the area under the ΔF/F curve during the stimulus period (0–3 s) for expanding stimuli. For non-expanding stimuli, a window of 1–3 s was used to exclude on-transients.

To estimate the contribution of each wasp component to the responses, we modelled each fly’s integrated responses to five stimuli (body + legs + antennae, body + legs, body + antennae, antennae, body) as *y* = *Xw*, where *X* is a binary matrix indicating the presence of components in each stimulus and *w ≥* 0 is a vector of weights fit by non-negative least squares. Each weight was normalized to the sum of weights across components to estimate each component’s share of the total drive. Shares were then averaged across flies.

### Connectomics analysis

#### Datasets

Analyses used either the male CNS connectome v1.0^30^ or FAFB connectome v783^31^. All main figure analyses were performed on the male CNS; Extended Data figures are labelled as FAFB where applicable.

#### LC-LC connectivity

We included all visual projection neuron types whose names begin with LC, LPLC, or LLPC (types annotated as unclear were excluded). Synapses outside of the lobula ROIs were excluded (LCs make nearly zero synapses with one another outside of the lobula). Within-type connections were excluded.

#### Trophic analysis

We computed trophic levels and trophic coherence for the weighted, directed, type-level LC network ^54^. With *W_ij_* as the synapse count from type *i* to type *j*, *u* = *w*^in^ + *w*^out^, and *v* = *w*^in^ *− w*^out^, trophic levels *h* solve Λ*h* = *v*, where Λ = diag(*u*) *− W − W* ^T^. Levels were then shifted such that the lowest node was positioned at 0. Trophic incoherence was defined as *F*_0_ = ∑*W_ij_*(*h_j_ − h_i_ −* 1)^2^*/*∑*W_ij_*, and trophic coherence was defined as 1 *−F*_0_. Networks that are more directed, with more or stronger feedforward edges, will have trophic coherence values closer to 1. A null distribution was obtained by generating 1,000 shuffled networks with degree-preserving swaps (no self-loops or duplicate edges), randomly permuting synapse counts across the rewired edges, and recomputing trophic levels and coherence for each shuffled network. We defined synapses within a trophic network as feedforward if the postsynaptic node was positioned at a higher trophic level than the presynaptic node.

#### LC-to-DN connectivity

We defined DNs as any neuron belonging to the superclass “descending_neuron”. For each of LC9, LC15, LC21, and LC25 we summed direct synapses from the LC type onto every downstream DN type and divided by the number of neurons of that DN type.

#### Indirect pathways to DNp09

Neurons mediating indirect input from LC15, LC21, and LC25 to DNp09 were identified by building a graph of all neurons on two-hop paths. With *J* as the neuron-level weight matrix and *x* as an indicator vector for each source LC type, we computed the downstream influence *V* = *x* + *xJ* + *xJ* ^2^. We removed each intermediate cell type in turn by zeroing its rows and columns in *J* and attributed the lost influence to that intermediate type.

#### Receptive field mapping

We mapped lobula columns to visual space using an eye map ^26^ of the male CNS (A. Zhao and M. Reiser, personal communication), which assigns a viewing direction for each ommatidium in hexagonal coordinates (*p*, *q*). For each LC type, we summed the input synapses across all its neurons within each lobula column. These synapses were then assigned to the corresponding (*p*,*q*) ommatidium in visual space. In Fig. 5g, both the ommatidial viewing angles and synapse counts were averaged across the left and right eyes/hemispheres. The receptive field that DNp09 inherits from each type was estimated by multiplying each LC neuron’s column-wise input synapses by the number of synapses that neuron makes onto DNp09, summed across neurons. We then averaged the two hemispheres and normalized the map to 100% (Fig. 5h). Azimuthal profiles are weighted histograms of column azimuths smoothed with a Gaussian kernel (*σ* = 10°) and normalized to their peak. The total connectomic prediction of DNp09’s receptive field (orange dashed trace in Fig. 5l) is the sum of the DNp09-weighted profiles of LC9, LC31a, and LC31b, each scaled by that type’s total weight onto DNp09.

To estimate the receptive field that DNp09 inherits from LC15, LC21, and LC25 through LC9, LC31a, and LC31b, we computed a path weight for each neuron *i* of the source type as ∑*_k_ w_ik_ w_k,_*_DNp09_, where *k* ranges over LC9, LC31a and LC31b neurons in the same hemisphere and *w* are synapse counts. Each source neuron’s column-wise input synapse counts were multiplied by its path weights, summed across neurons within a type, averaged across hemispheres, and then normalized to 100%.

#### Comparison of connectomic predictions with turning behaviour

Behavioural turning rates and connectomic predictions were compared by fitting a binomial generalized linear model (logit link) of turn counts per azimuth bin (turns out of frames, left and right turns mapped onto signed azimuth) with the peak-normalized connectomic curve as predictor. The slope’s 95% confidence interval and *P* value were derived from standard errors clustered by fly (t distribution, *n −* 1 degrees of freedom). The 95% confidence band for the turning-rate curve (Fig. 5l) and the 95% confidence interval for the Pearson correlation between the turning-rate and connectomic curves were obtained from 2,000 bootstrap resamples of flies. Predictive performance (Extended Data Fig. 9b) was quantified as held-out deviance explained relative to a constant turning probability, in five-fold cross-validation over flies repeated 20 times. Differences in deviance explained between the connectomic prediction and each alternative were estimated from 1,000 paired bootstrap resamples of flies.

## Acknowledgements

We thank A. Zhao and M.B. Reiser for sharing male CNS eyemaps; B. Moon for insect illustrations used in the manuscript; L.F. Abbott, J. Clemens, S. Govind, D.J.C. Kronauer, J.L. Zung, the Smithsonian Tropical Research Institute (STRI) community, and members of the Axel, Behnia, and Card labs for discussions; M.J. Hughes and L.S. Kors for technical support; and M. D’Arduini, G.G. Glinka, M. Gutierrez, and A. Nemes for general laboratory support. Stocks from the Bloomington Drosophila Stock Center (NIH P40OD018537) and National Drosophila Species Stock Center (NDSSC, Cornell University) were used in this study.

## Author information

### Contributions

Authors are listed alphabetically by surname. Conceptualization: R.A., G.M.C., K.M.C., S.D.; Methodology: K.M.C., S.D.; Investigation: K.M.C., S.D., M.F.; Resources: V.M.D.; Data curation, Formal analysis: K.M.C., S.D.; Visualization: G.M.C., K.M.C., S.D.; Writing – original draft: R.A., G.M.C., K.M.C., S.D.; Writing – review & editing: R.A., R.B., G.M.C., K.M.C., S.D.; Supervision, Project administration: R.A., R.B., G.M.C., K.M.C.; Funding acquisition: R.A., R.B., G.M.C., K.M.C., S.D.

### Corresponding author

Correspondence to Kevin M. Cury.

## Extended Data Figures and Tables

**Extended Data Figure 1.**
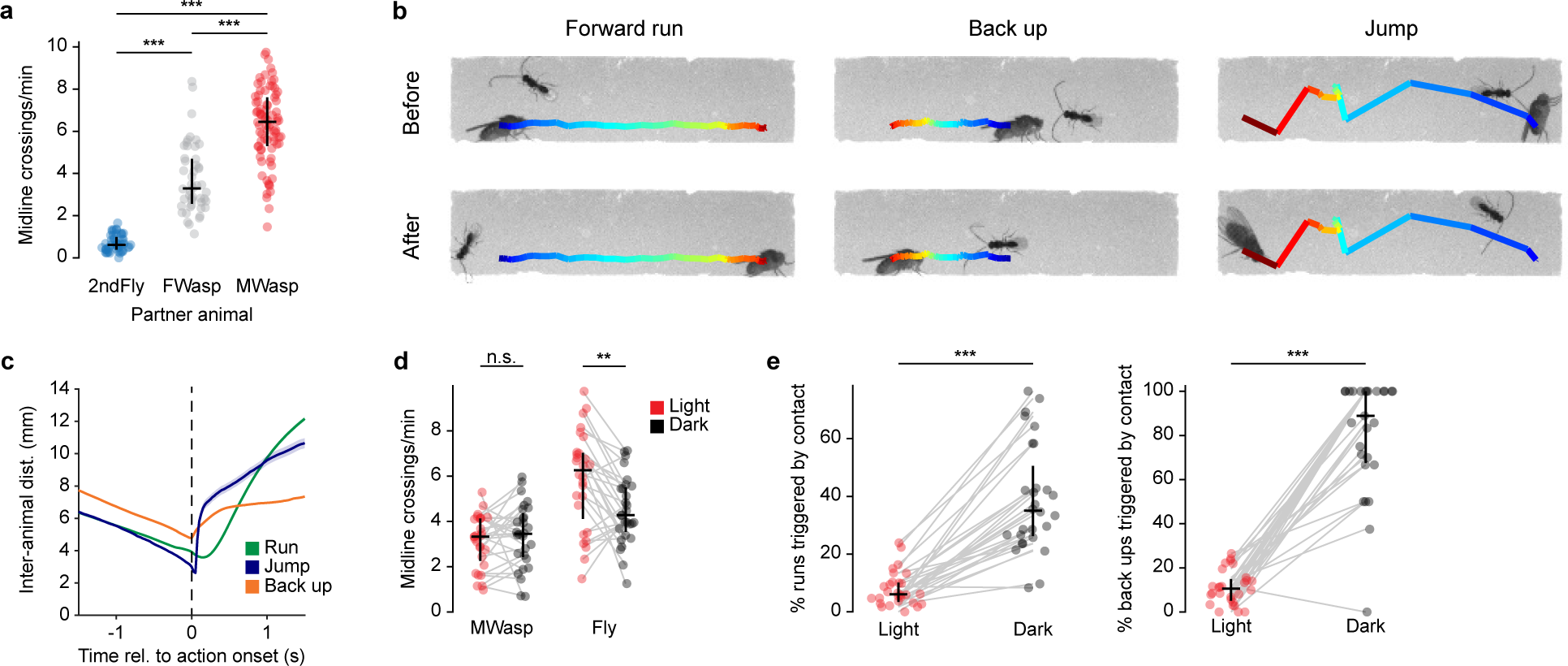
**a,** The rate of fly midline crossings across all three pairings. Here and in d and e: points, individual pairings; horizontal lines, median; vertical line, interquartile range (25^th^–75^th^ percentile). *n* (pairs): 2ndFly, 52; FWasp, 48; MWasp, 82. One-way ANOVA: *F* (2, 179) = 235.77, *P* = 6.96 × 10^−51^. Tukey–Kramer post hoc test: 2ndFly vs. FWasp, *P* = 1.14 × 10^−19^, and 2ndFly vs. MWasp, *P* = 2.06 × 10^−51^; FWasp vs. MWasp, *P* = 1.73 × 10^−22^. Data are from the same animal pairs as in Figure 1b–d. **b,** Representative evasive actions performed by flies paired with male wasps. Screenshots at the onset (top) and offset (bottom) of three distinct evasive actions, with the intermediate trajectory of the fly’s central midpoint colour-coded. Before, time = 0 s; After, time = 1.475 s (Forward run), 1.1125 s (Back up), 0.525 s (Jump). See also Supplementary Videos 4–6. **c,** Inter-animal distance in the ±1.5 s surrounding each evasive action (action onset at time = 0) performed by flies paired with male wasps; line and shading, mean ± s.e.m. *n* (events; from 82 pairs): Forward run, 1,457; Back up, 1,877; Jump, 389. Only scoring subset of actions falling within iso-contour regions drawn at 50% of the peak event density. Only events occurring while the fly was on the chamber sidewalls were considered here. **d,** The rate of midline crossings for both animals for pairings of a female fly with a male wasp, measured in paired lights-on and dark sessions.. Here and in e, *n* = 28 pairs. Paired t-test on light versus dark: MWasp, N.S.; female fly, *P* = 1.19 × 10^−3^. Data are from the same animal pairs as in Figure 1e. **e,** Fraction of forward running (left panel) and backing-up (right panel) events occurring within 50 ms of contact for flies paired with male wasps, measured in paired lights-on and dark sessions.. Paired t-test on light versus dark: Forward run, *P* = 2.19 × 10^−10^; Back up, *P* = 8.44 × 10^−13^.

**Extended Data Figure 2.**
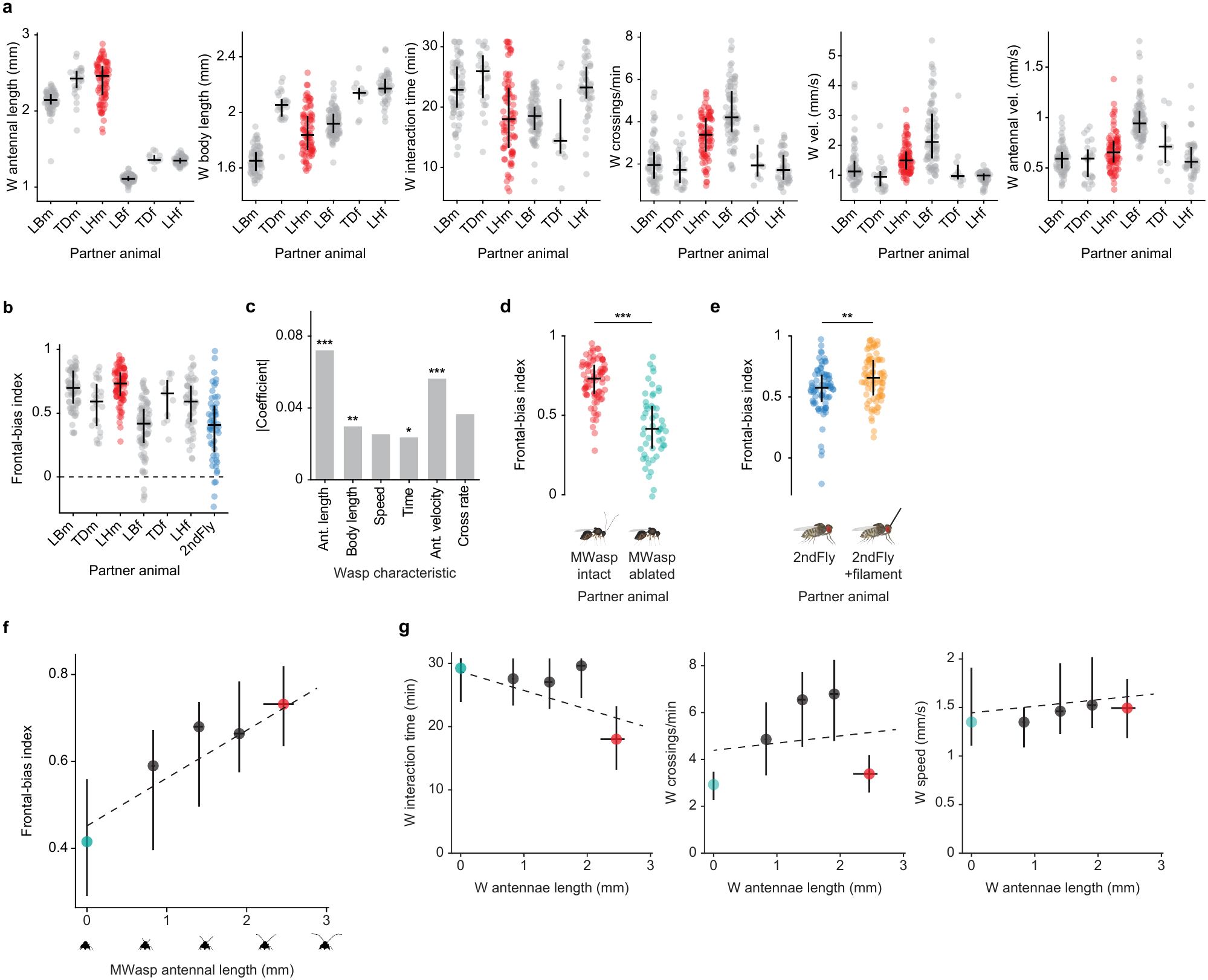
**a,** Morphological and behavioural characteristics of wasps for the males and females of three *Drosophila*-targeting parasitoid wasp species: LB, *Leptopilina boulardi*; TD, *Trichopria drosophilae*; LH, *Leptopilina heterotoma*. Left to right: antennal length, body length, time spent in chamber, midline-crossing rate, speed at midline crossing, antennal tip velocity. Here and in b and c: male *L. boulardi*, *n* = 65 pairs; male *T. drosophilae*, *n* = 25 pairs; male *L. heterotoma*, *n* = 82 pairs; female *L. boulardi*, *n* = 80 pairs, female *T. drosophilae*, *n* = 11 pairs; female *L. heterotoma*, *n* = 48 pairs; male *D. micromelanica*, *n* = 52 pairs. Here and in b, d, e, and g: points, individual pairings; horizontal lines, median; vertical line, interquartile range (25^th^–75^th^ percentile). Data are from the same animal pairs as in Figure 2a, b. **b,** Frontal-bias index of flies paired with males or females of three wasp species and a male fly of a different species. **c,** Coefficient magnitudes from a multiple linear regression model with six morphological and behavioural predictors of fly frontal-bias index. Antennal length, *P* = 1.33 × 10^−8^; body length, *P* = 0.00875; midline-crossing rate, *P* = 0.0591; interaction time, *P* = 0.0488; antennal velocity, *P* = 0.000560; speed at midline crossing, *P* = 0.0943. Antennal length and speed at midline crossing had positive coefficients; all others, negative. The model explained 64% of the variance in frontal-bias index across group means. **d,** Frontal-bias index of flies paired with individual male wasps (*L. heterotoma*) with intact (MWasp intact) or fully removed antennae (MWasp ablated). *n* (pairs): MWasp intact, 82; MWasp ablated, 50. Two-sided two-sample t-test: *P* = 9.01 × 10^−17^. Data are from the same animal pairs as in Figure 2c. **e,** Frontal-bias index of flies paired with individual heterospecific flies without (2ndFly) or with (2ndFly+filament) a synthetic filament affixed to their heads. *n* (pairs): 2ndFly, 75; 2ndFly+filament, 80. Two-sided two-sample t-test: *P* = 0.00233. Data are from the same animal pairs as in Figure 2d. **f,** Frontal-bias index of flies versus wasp antennal length for pairings with male wasps whose antennae were cut to four distinct lengths, or left intact. Here and in g, *n* (pairs): 0 segments, 50; 3 segments, 61; 6 segments, 60; 9 segments, 58; 13 segments (intact), 82. Linear regression: intercept = 0.452, slope = 0.110, *R*^2^ = 0.213, *P* = 9.08 × 10^−18^ (slope). Data are from the same animal pairs as in Figure 2e. **g,** Behavioural characteristics of wasps whose antennae were cut to four distinct lengths, or left intact. From left to right: time spent in chamber, midline-crossing rate, speed at midline crossing. Linear regression: antennal length vs. time in chamber, intercept = 28.697, slope = −2.980, *R*^2^ = 0.156, *P* = 4.65 × 10^−13^ (slope); antennal length vs. midline-crossing rate, intercept = 4.387, slope = 0.308, *R*^2^ = 0.010, *P* = 0.0782; antennal length vs. speed at midline crossing, intercept = 1.445, slope = 0.067, *R*^2^ = 0.011, *P* = 0.0655.

**Extended Data Figure 3.**
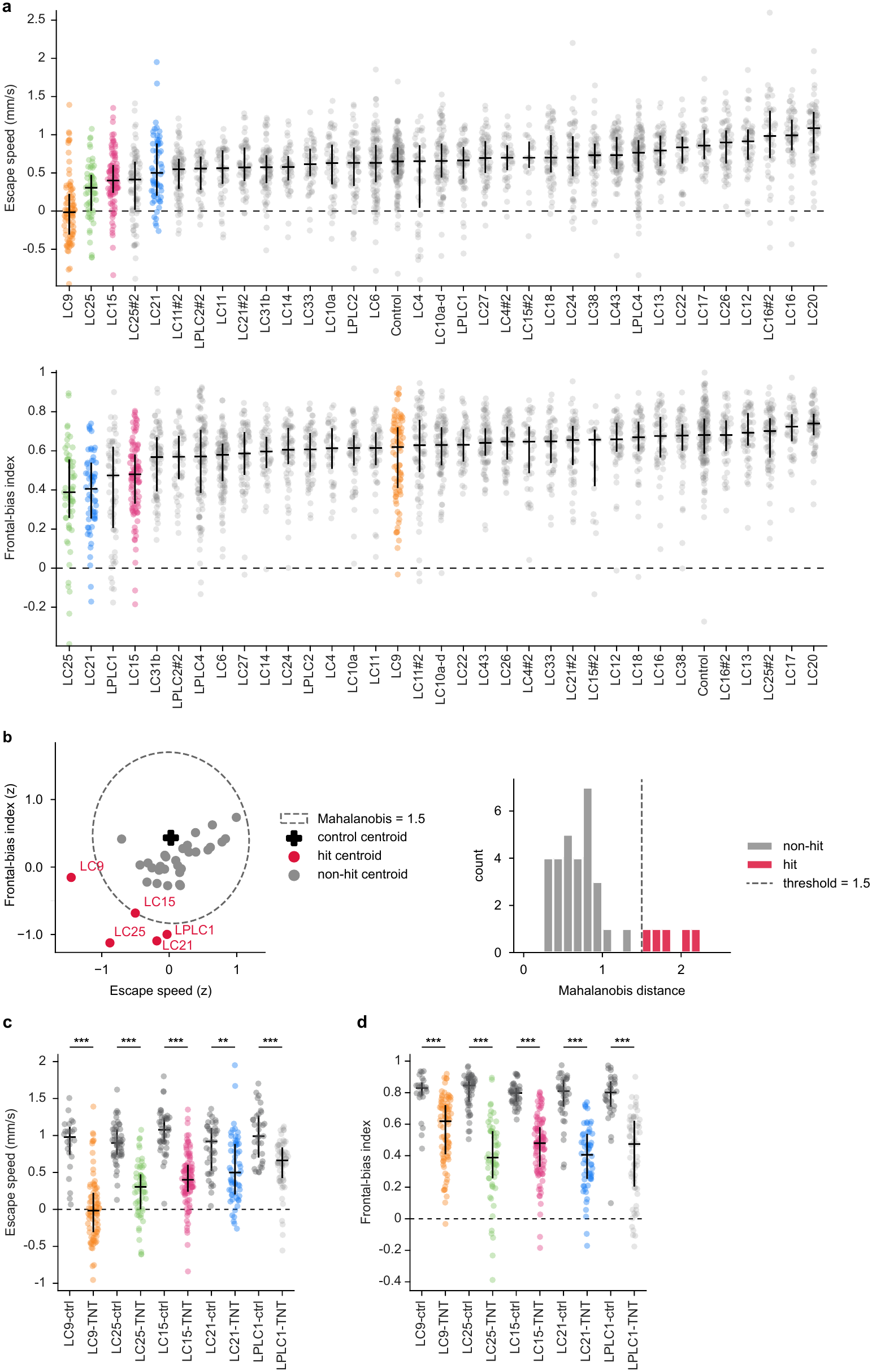
**a,** Top, escape speed for each silenced LC type and the empty split-GAL4 control. Here and in Extended Data Fig. 3b–d: points, individual pairings; horizontal lines, median; vertical line, interquartile range (25^th^–75^th^ percentile). Bottom, frontal-bias index for each silenced LC type and the empty split-GAL4 control. Here and in b, *n* (pairs; split-GAL4>TNT): Control, 145; LC4, 44; LC4 #2, 42; LC6, 106; LC9, 82; LC10a, 58; LC10a-d, 89; LC11, 53; LC11 #2, 81; LC12, 54; LC13, 60; LC14, 52; LC15, 97; LC15 #2, 23; LC16, 57; LC16 #2, 50; LC17, 45; LC18, 62; LC20, 65; LC21, 58; LC21 #2, 59; LC22, 50; LC24, 65; LC25, 57; LC25 #2, 79; LC26, 63; LC27, 69; LC31b, 77; LC33, 54; LC38, 57; LC43, 82; LPLC1, 63; LPLC2, 65; LPLC2 #2, 50; LPLC4, 110. Data are from the same animal pairs as in Figure 3b. **b,** Distance criteria for identifying LC types whose silencing altered defensive behaviour relative to control. Left, centroids of each split-GAL4>TNT group in the space defined by escape speed and frontal-bias index, with both metrics z-scored; cross, control centroid (empty split-GAL4>TNT); dashed ellipse, Mahalanobis distance of 1.5 from control; red, groups exceeding this distance threshold; grey, all other groups. Right, distribution of Mahalanobis distances from control across LC split-GAL4>TNT groups. Dashed line, distance threshold of 1.5. **c,** Escape speed for five LC types whose silencing altered defensive behaviour relative to control (as identified in b), each compared with its corresponding GAL4-only genetic control (LC split-GAL4 crossed to Canton-S). Here and in d, *n*: LC9>TNT, 82; LC9-ctrl, 24; LC25>TNT, 57; LC25-ctrl, 51; LC15>TNT, 97; LC15-ctrl, 44; LC21>TNT, 58; LC21-ctrl, 39; LPLC1>TNT, 63; LPLC1-ctrl, 33. Two-sided two-sample t-test (LC>TNT versus LC-ctrl): LC9, *P* = 4.86 × 10^−14^; LC25, *P* = 4.67 × 10^−17^; LC15, *P* = 7.95 × 10^−19^; LC21, *P* = 0.00136; LPLC1, *P* = 3.61 × 10^−7^. Same LC split-GAL4>TNT data depicted in a. **d,** Frontal-bias index for five LC types whose silencing altered defensive behaviour relative to control (as identified in b), each compared with its corresponding GAL4-only genetic control (LC split-GAL4 crossed to Canton-S). Two-sided two-sample t-test (LC>TNT versus LC-ctrl): LC9, *P* = 1.63 × 10^−6^; LC25, *P* = 2.32 × 10^−19^; LC15, *P* = 1.67 × 10^−20^; LC21, *P* = 1.10 × 10^−16^; LPLC1, *P* = 1.49 × 10^−10^. Same LC split-GAL4>TNT data depicted in a.

**Extended Data Figure 4.**
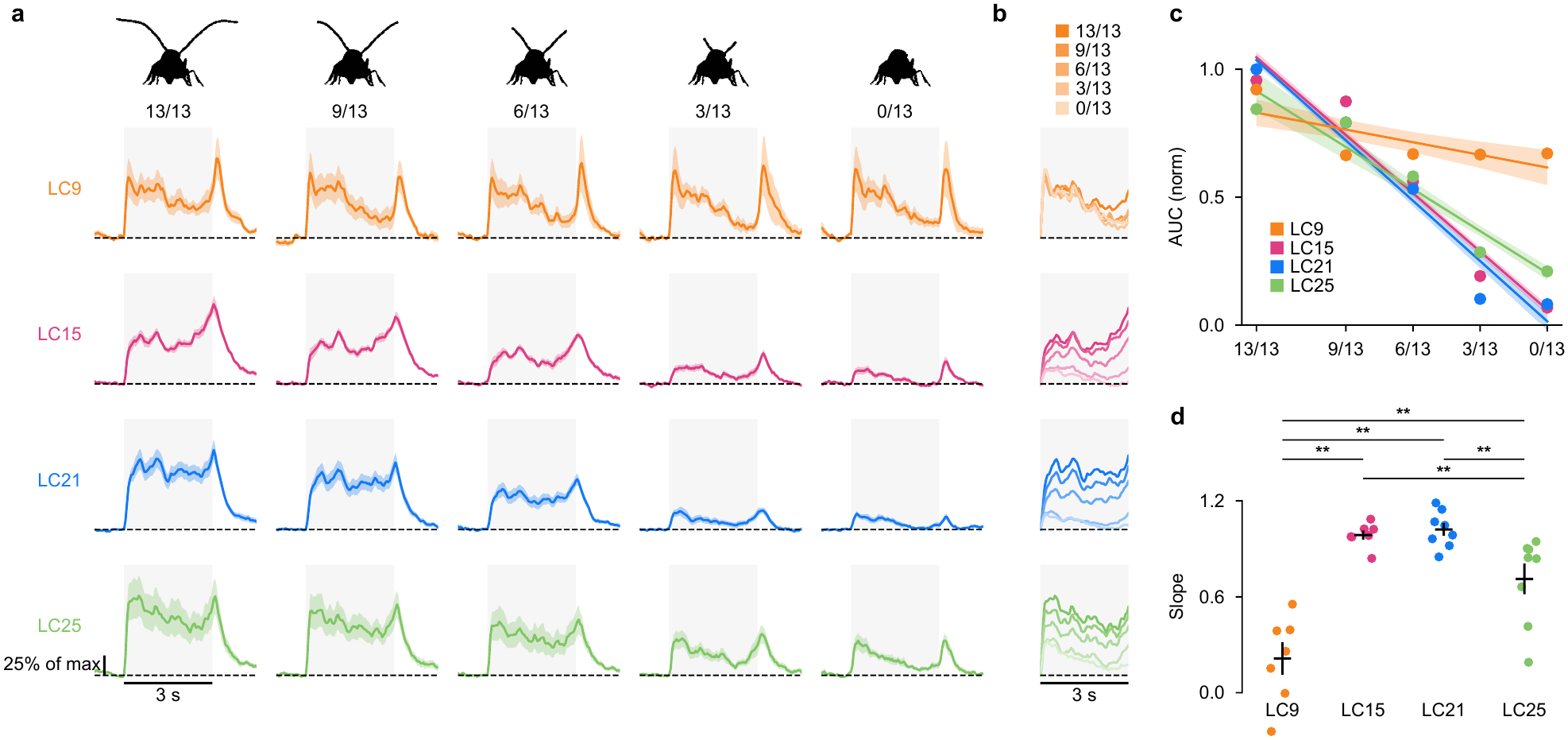
**a,** Mean calcium responses of LC9, LC15, LC21, and LC25 (rows) to non-expanding movies of walking wasps whose antennae were truncated to 13/13, 9/13, 6/13, 3/13, or 0/13 of their full length (columns). For each cell type, responses are normalized to the peak response to the intact (13/13) stimulus. Grey shading, stimulus period (0–3 s); dashed line ΔF/F = 0; line and shading, mean ± s.e.m. *n* (flies): LC9, 7; LC15, 7; LC21, 8; LC25, 8. **b,** Mean responses during the stimulus period (0–3 s) from a overlaid for each cell type. Shading indicates antennal length, from 13/13 segments (dark) to 0/13 segments (light). **c,** Integrated responses (area under the curve, 1–3 s after stimulus onset) as a function of antennal length. For each fly, responses were normalized to the maximum across the five antennal lengths. Points, mean across flies; lines and shading, mean ± s.e.m. of linear fits to individual flies. Same flies as in a. **d,** Slopes of the per-fly linear fits in c. Points, individual flies; crosses, mean ± s.e.m. Kruskal–Wallis test, *H* = 21.4, *P* = 8.89 × 10^−5^; two-sided Mann–Whitney *U* tests with Bonferroni correction: LC9 vs. LC15, *P* = 5.83 × 10^−4^; LC9 vs. LC21, *P* = 3.11 × 10^−4^; LC9 vs. LC25, *P* = 5.91 × 10^−3^; LC15 vs. LC21, *P* = 0.613; LC15 vs. LC25, *P* = 3.73 × 10^−3^; LC21 vs. LC25, *P* = 1.86 × 10^−3^.

**Extended Data Figure 5.**
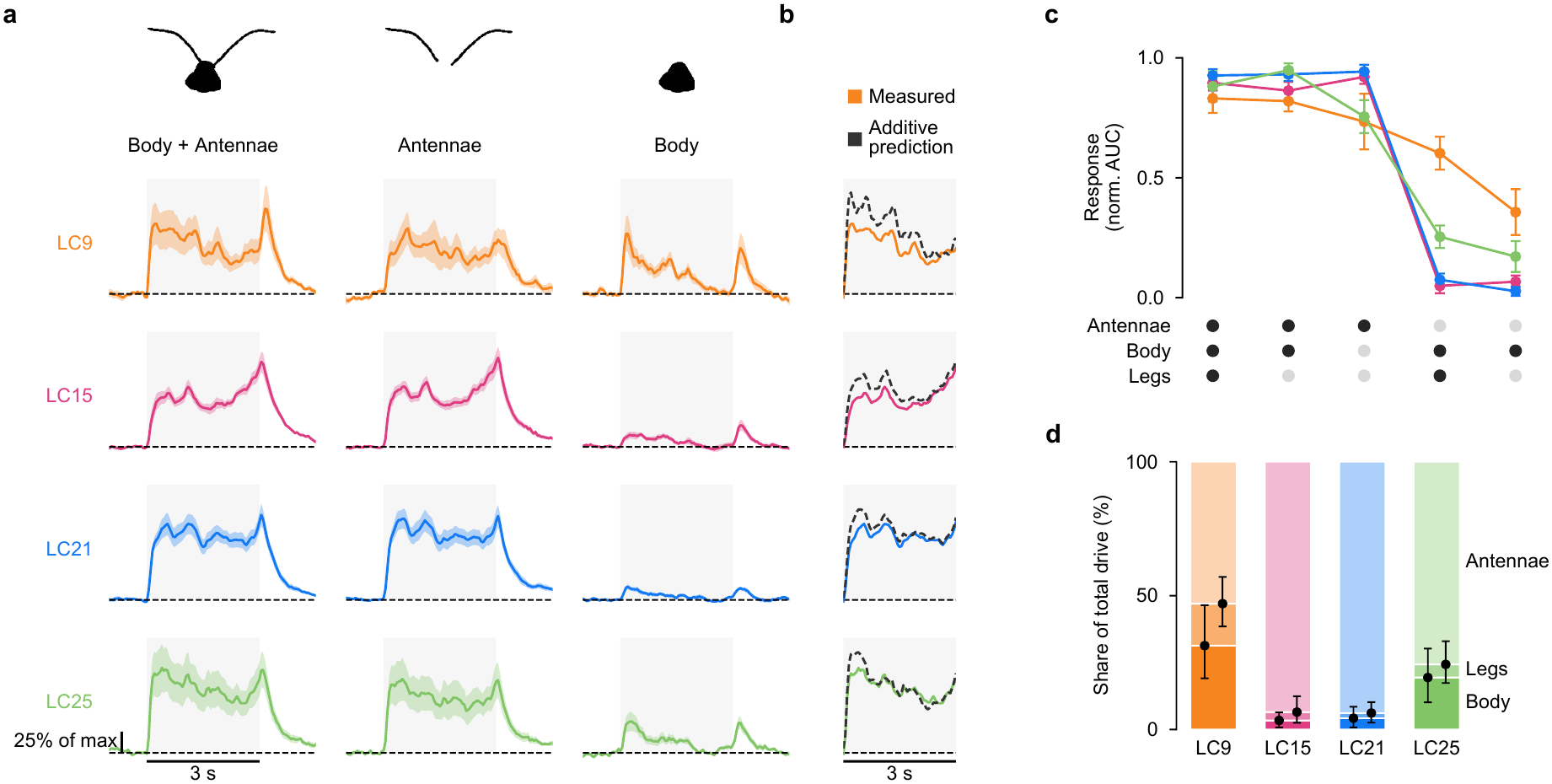
**a,** Mean calcium responses of LC9, LC15, LC21, and LC25 (rows) to non-expanding movies of walking wasps showing the body and antennae together (left), the antennae alone (middle), or the body alone (right). For each cell type, responses are normalized to the peak response to body + antennae stimulus. Grey shading, stimulus period (0–3 s); dashed line ΔF/F = 0; line and shading, mean ± s.e.m. *n* (flies): LC9, 7; LC15, 7; LC21, 8; LC25, 8. **b,** Measured response to the body + antennae stimulus (coloured; from column one of a) compared with the additive prediction (dashed black), computed for each fly as the sum of its responses to the antennae-alone and body-alone stimuli. **c,** Integrated responses (area under the curve, 1–3 s after stimulus onset) to five non-expanding stimuli containing different combinations of wasp components (dot matrix below; black, component present). The body + legs + antennae and body + legs stimuli are the 13/13 and 0/13 stimuli of Extended Data Fig. 4a, respectively; the body + antennae, antennae-only and body-only stimuli are those shown in a. For each fly, responses were normalized to the maximum across the five stimuli. Points and error bars, mean ± s.e.m. across flies. **d,** Contribution of each wasp component to the integrated response to non-expanding stimuli for each LC type. For each fly, integrated responses to the five stimuli in c were fit with a non-negative additive model, and the fitted weights were expressed as shares of the total (see Methods). Stacked bars, mean share attributed to the body (dark), legs (intermediate), and antennae (light). Points and error bars, mean and 95% confidence interval of boundaries. Median *R*^2^ of the additive fit across flies: LC9, 0.399; LC15, 0.971; LC21, 0.986; LC25, 0.968.

**Extended Data Figure 6.**
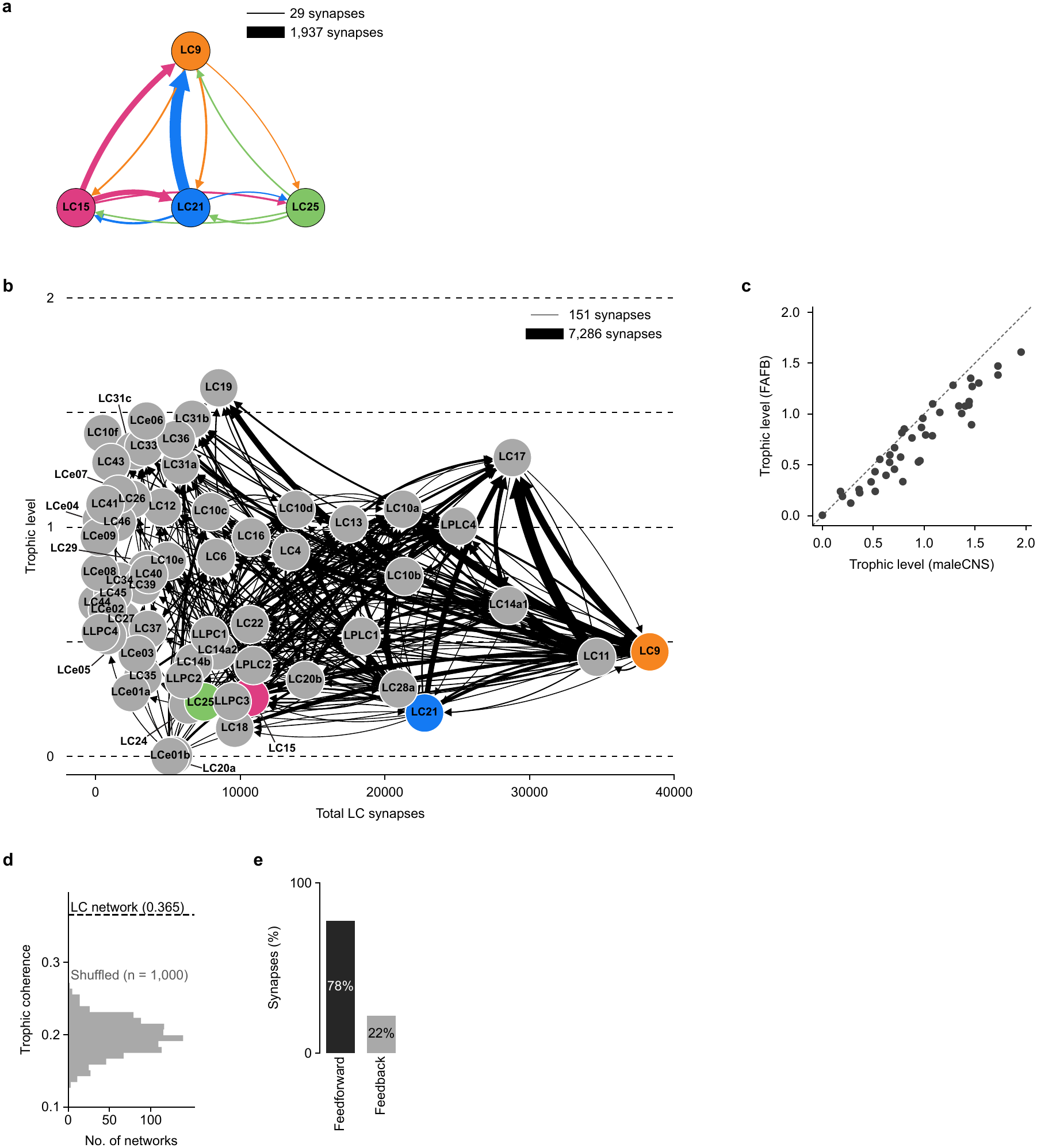
**a,** Synaptic connectivity among LC9, LC15, LC21 and LC25 within the lobula of the FAFB connectome, both hemispheres. Arrows point from the presynaptic to the postsynaptic type and are coloured by the presynaptic type; arrow width is proportional to synapse count (range, 29–1,937 synapses). Within-type connections are not shown. Compare with Figure 4e. **b,** Trophic analysis of the FAFB LC-to-LC network (62 LC, LPLC and LLPC types; lobula synapses only), plotted as in Fig. 4f. Each node is an LC type, positioned by its trophic level (y axis) and its total LC-to-LC synapses (input + output; x axis). Edge width is proportional to synapse count, and only connections of more than 150 synapses are shown. LC9, LC15, LC21 and LC25 are coloured. **c,** Trophic levels of the 41 LC types present in both networks, FAFB versus male CNS. Pearson *r* = 0.95. **d,** Trophic coherence of the FAFB LC-to-LC network (dashed line, 0.365) compared with 1,000 shuffled networks generated by degree-preserving rewiring followed by permutation of synapse counts (grey histogram; 0.196 ± 0.025, mean ± s.d.). *z* = 6.9; empirical *P* < 0.001 (two-sided). **e,** Percentage of FAFB LC-to-LC synapses directed up the trophic hierarchy (feedforward; 78%) or down it (feedback; 22%), weighted by synapse count.

**Extended Data Figure 7.**
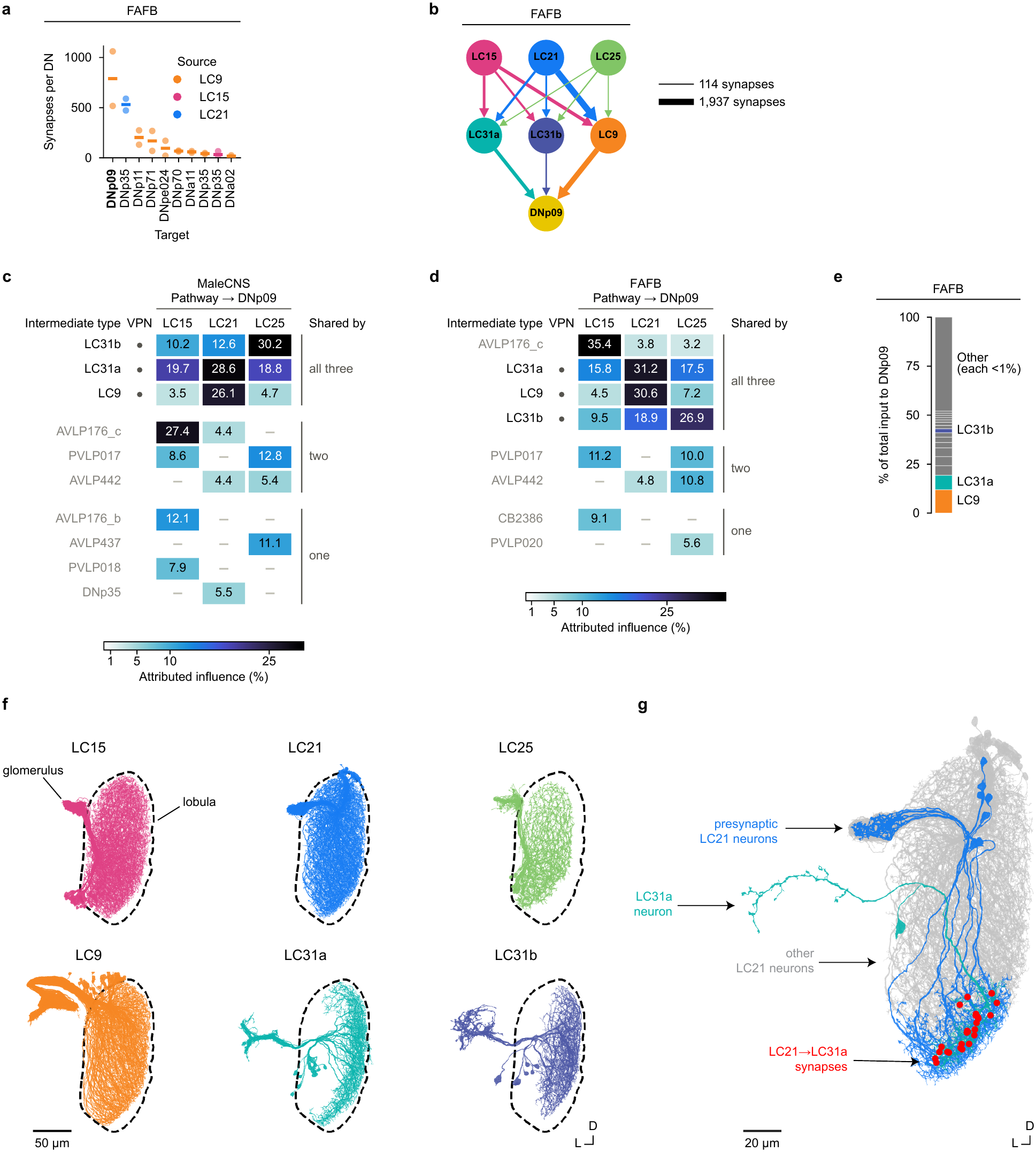
**a,** Input synapses per descending neuron for the strongest direct connections from LC9, LC15, LC21 and LC25 onto DN types in FAFB, plotted as in Figure 5b. Points, individual DNs; lines, mean. **b,** Connectivity from LC15, LC21 and LC25 onto LC9, LC31a and LC31b, and from these onto DNp09, in FAFB, plotted as in Figure 5c. Edge width is proportional to synapse count. **c,** Intermediate cell types that carry indirect input from LC15, LC21 and LC25 to DNp09 in the male CNS connectome. Each cell is the percentage of two-hop influence from the source LC type (column) onto DNp09 that is lost when the intermediate type (row) is removed, averaged over the two DNp09 neurons (see Methods). Rows are the union of the ten strongest intermediates for each source type, grouped by the number of source types they share. Only values *≥* 1% are shown, for rows whose total is *≥* 5%. LC9, LC31a and LC31b are highlighted, and visual projection neuron types are marked. **d,** Same analysis as c for FAFB. **e,** DNp09 inputs in FAFB, ranked by the proportion of input synapses they contribute (13,216 synapses across both DNp09 neurons). LC9, 11.9%; LC31a, 7.4%; LC31b, 2.0%. Any inputs that individually contribute <1% of synapses are pooled as Other. Compare with Figure 5e. **f,** EM reconstructions of the right-hemisphere populations of LC15 (65 neurons), LC21 (78), LC25 (27), LC9 (115), LC31a (16) and LC31b (6) in the male CNS connectome. Scale bar, 50 µm. **g,** A single LC31a neuron (male CNS, right hemisphere; body ID 25708) and the seven LC21 neurons presynaptic to it (27 synapses; synapse locations marked). The remaining 71 right-hemisphere LC21 neurons are shown in grey. Scale bar, 20 µm.

**Extended Data Figure 8.**
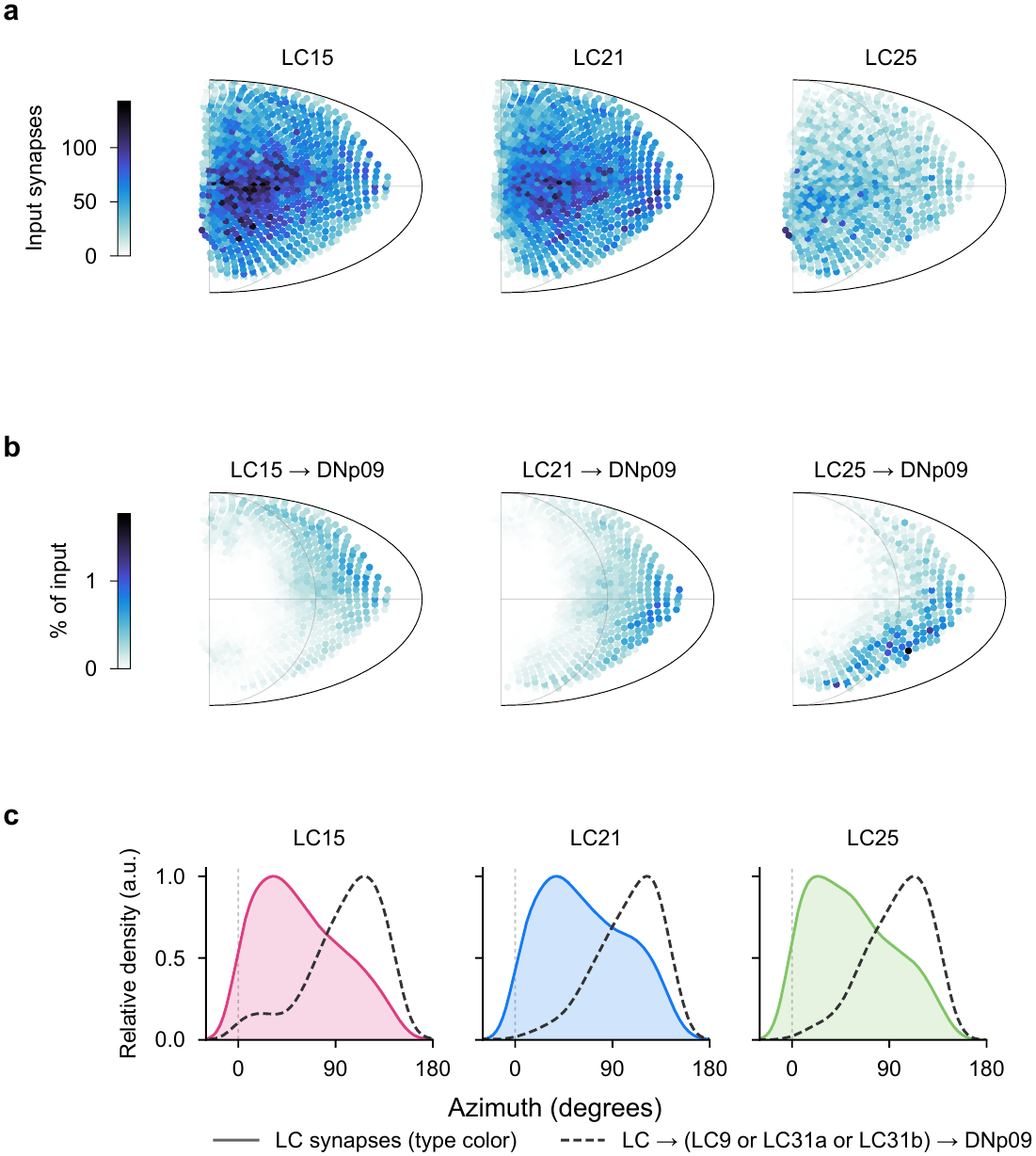
**a,** Predicted receptive fields of the LC15, LC21 and LC25 populations, shown as Mollweide projections as in Fig. 5g. Each point is an ommatidium positioned at its viewing direction (Mollweide projection) and coloured by the number of input synapses from the corresponding lobula column. The map for the left eye was mirrored about the 0°–180°azimuthal meridian and averaged with the right eye map. **b,** Receptive fields that DNp09 inherits from LC15, LC21 and LC25 through the two-hop pathway via LC9, LC31a and LC31b, shown as Mollweide projections as in Fig. 5h. Each point is coloured by the percentage of two-hop input contributed by the corresponding lobula column (see Methods). The map for the left eye was mirrored about the 0°–180°azimuthal meridian and averaged with the right eye map. **c,** Azimuthal distributions of the receptive field of each LC type (solid, coloured) and of its two-hop contribution to DNp09 (dashed). Each curve is normalized to its peak; Gaussian smoothing, *σ* = 10°.

**Extended Data Figure 9.**
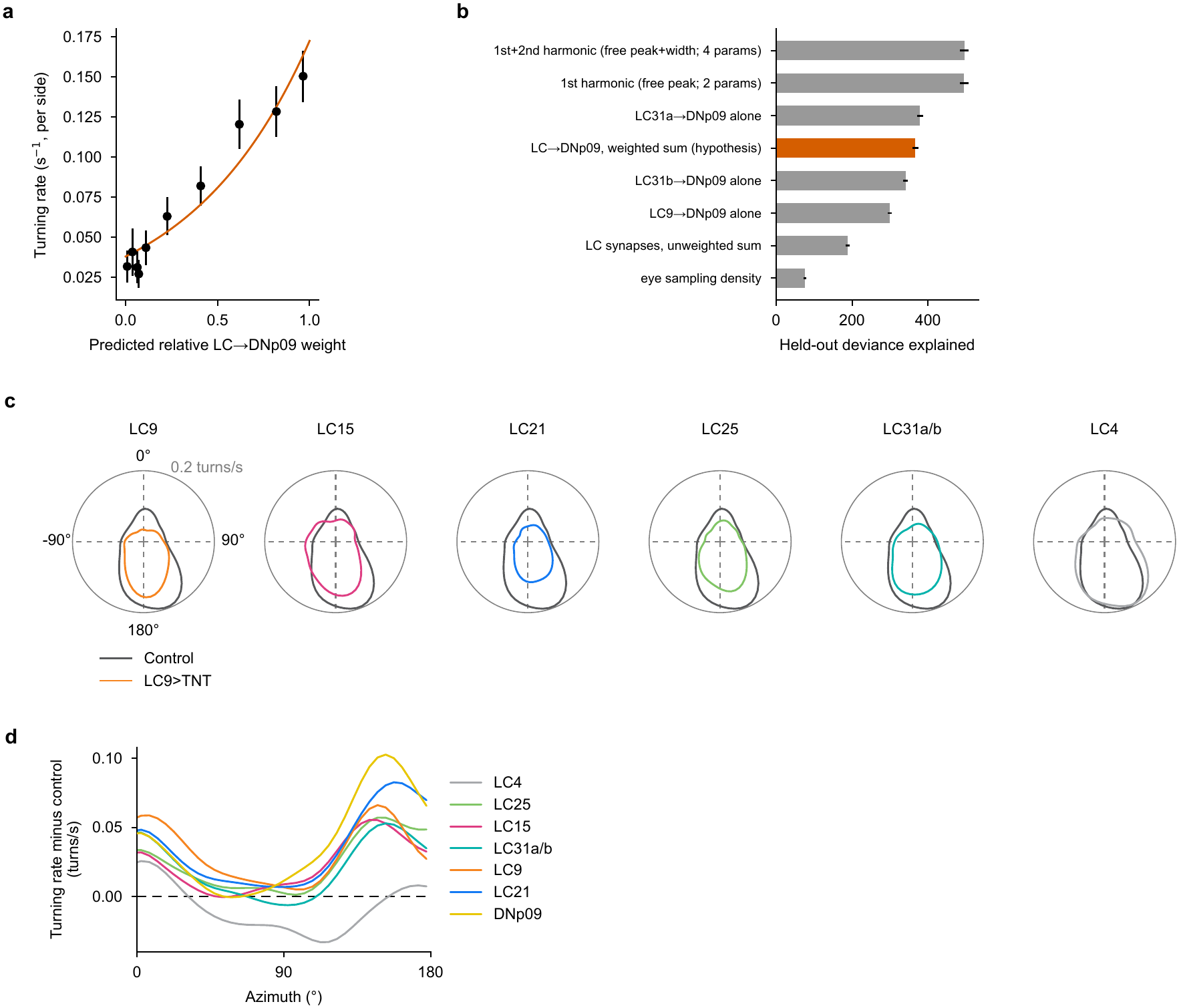
**a,** Turning rate per side (mean, wild-type flies paired with male wasps) in deciles of the predicted relative LC*→*DNp09 weight, with the binomial GLM fit (error bars, 95% CI). *n* = 1,574 turns from 57 flies. **b,** Held-out deviance explained (fivefold cross-validation over flies, 20 repeats; error bars, s.d.) by the connectomic prediction (LC9 + LC31a + LC31b *→* DNp09, weighted sum) and by alternative predictors. Alternatives: each LC type alone, the unweighted LC9 + LC31a + LC31b synapse density, and the ommatidial sampling density. Ceilings are one- and two-harmonic tuning curves fit directly to the behaviour. The connectomic prediction reached 81% of the two-harmonic ceiling and outperformed the unweighted synapse density and the eye sampling density. It was not distinguishable from LC31a alone. **c,** Rate of turns toward the wasp as a function of wasp azimuth for flies expressing TNT in LC9, LC15, LC21, LC25, LC31a/b or LC4 (coloured), each overlaid with the empty split-GAL4 control (grey), paired with male wasps in a circular chamber. Plotted as in Fig. 5m: 0°, wasp directly in front of the fly; left turns mirrored. Turns: control, 1,583; LC9, 1,020; LC15, 1,544; LC21, 1,212; LC25, 1,287; LC31a/b, 1,250; LC4, 2,998. **d,** Difference in toward-turn rate between controls and each silenced genotype (control minus silenced) as a function of wasp azimuth on the side of the turn. Positive values indicate reduced turning in silenced flies.

## Supplementary information

### Supplementary Video 1

Fly–fly interaction in a rectangular chamber. Representative video recording of the initial 4 min of the lights-on session of an interaction between a female *D. melanogaster* fly and a male heterospecific fly (*D. micromelanica*). The female *D. melanogaster* fly is lighter in shade and larger in size. Corresponding behavioural traces shown in Figure 1a, left column. Video playback speed is 10× real time.

### Supplementary Video 2

Fly–female wasp interaction in a rectangular chamber. Representative video recording of the initial 4 min of the lights-on session of an interaction between a female *D. melanogaster* fly and a female *L. heterotoma* parasitoid wasp. Corresponding behavioural traces shown in Figure 1a, middle column. Video playback speed is 10× real time.

### Supplementary Video 3

Fly–male wasp interaction in a rectangular chamber. Representative video recording of the initial 4 min of the lights-on session of an interaction between a female *D. melanogaster* fly and a male *L. heterotoma* parasitoid wasp. Corresponding behavioural traces shown in Figure 1a, right column. Video playback speed is 10× real time.

### Supplementary Video 4

Fly forward-running events observed during interactions with male wasps in a rectangular chamber. 20 representative videos of forward-running events by female *D. melanogaster* flies during interactions with male *L. heterotoma* parasitoid wasps. The red box outlining the video indicates the timing of each forward-running event, from onset to offset.

### Supplementary Video 5

Fly backing-up events observed during interactions with male wasps in a rectangular chamber. 20 representative videos of backing-up events by female *D. melanogaster* flies during interactions with male *L. heterotoma* parasitoid wasps. The red box outlining the video indicates the timing of each backing-up event, from onset to offset.

### Supplementary Video 6

Fly jumping events observed during interactions with male wasps in a rectangular chamber. 20 representative videos of jumping events by female *D. melanogaster* flies during interactions with male *L. heterotoma* parasitoid wasps. The red box outlining the video indicates the timing of each jumping event, from onset to offset.

### Supplementary Video 7

Fly–male wasp interaction in a circular chamber. Representative video recording of the initial 4 min of the lights-on session of an interaction between a female *D. melanogaster* fly and a male *L. heterotoma* parasitoid wasp. Corresponding behavioural traces shown in Figure 1g. Video playback speed is 10× real time.

### Supplementary Video 8

Representative turning events by a female *D. melanogaster* fly during interaction with a male *L. heterotoma* parasitoid wasp in a circular chamber. The red box outlining the video indicates the timing of each turning event, from onset to offset. Video playback speed is 4× real time.

### Supplementary Video 9

Fly–male wasp interactions in a rectangular chamber. Two representative video recordings each showing the initial 4 min of the lights-on session of an interaction between a female *D. melanogaster* fly and a male *L. heterotoma* parasitoid wasp with intact (MWasp intact; first video clip) or fully removed antennae (MWasp ablated; second video clip). Video playback speed is 10× real time.

### Supplementary Video 10

Fly–fly interactions in a rectangular chamber. Two representative video recordings each showing 6 min of an interaction between a female *D. melanogaster* fly and a male heterospecific fly (*D. micromelanica*) without (2ndFly; first video clip) or with (2ndFly+filament; second video clip) a synthetic filament affixed to its head. Video playback speed is 20× real time.

### Supplementary Video 11

Three movie clips (rows) of walking wasp silhouettes expanding to simulate approach whose antennae were full length (13/13) or truncated to 9/13, 6/13, 3/13, or 0/13 of their full length (columns).

### Supplementary Video 12

Three movie clips (rows) of non-expanding walking wasp silhouettes whose antennae were full length (13/13) or truncated to 9/13, 6/13, 3/13, or 0/13 of their full length (columns).

### Supplementary Video 13

Three movie clips (rows) of walking wasp silhouettes expanding to simulate approach in which the wasp was deconstructed and presented as various subsets of its components (columns). B, body; A, antennae; L, legs.

### Supplementary Video 14

Three movie clips (rows) of non-expanding walking wasp silhouettes in which the wasp was deconstructed and presented as various subsets of its components (columns). B, body; A, antennae; L, legs.

**Supplementary Table 1.**
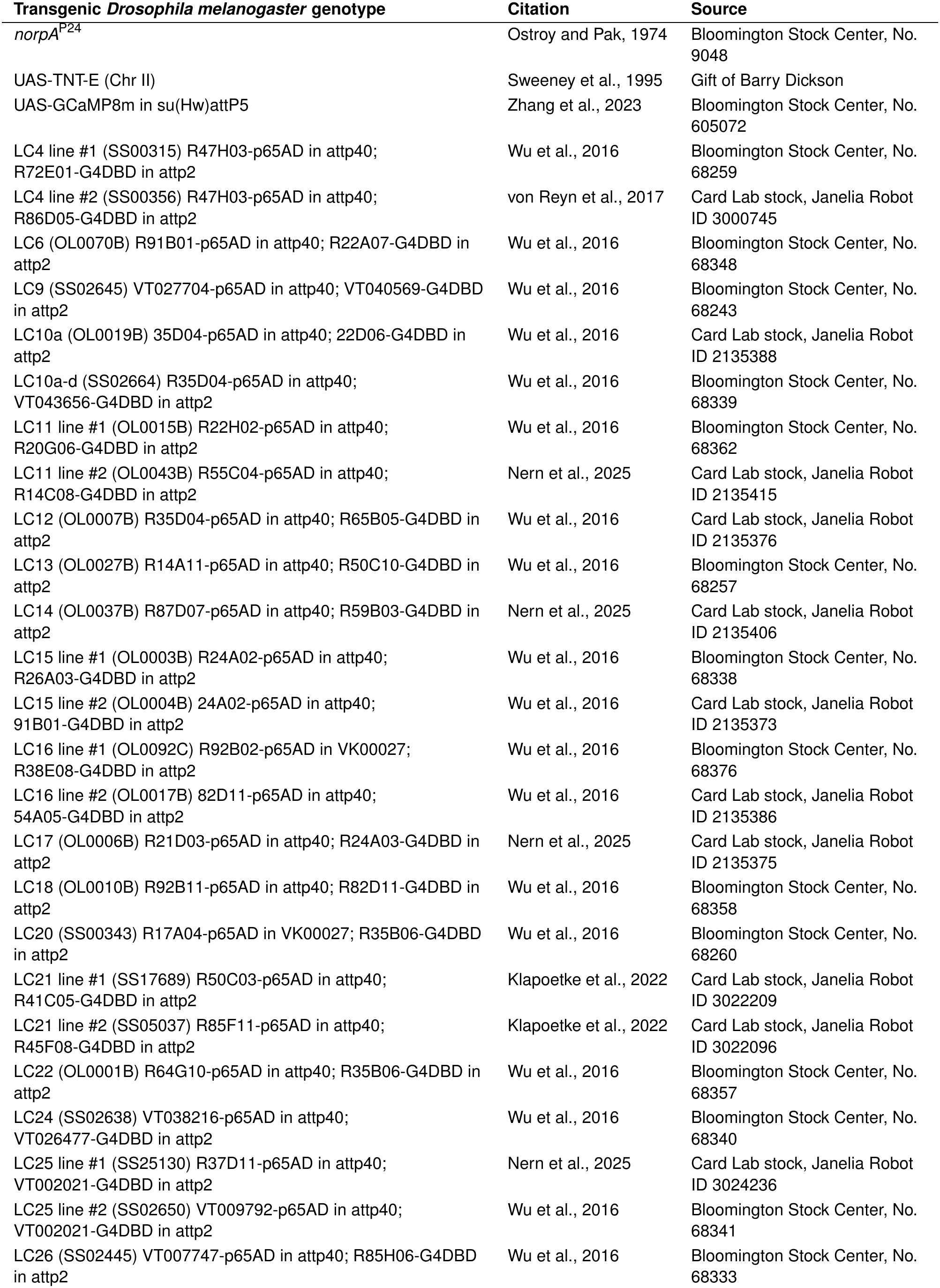
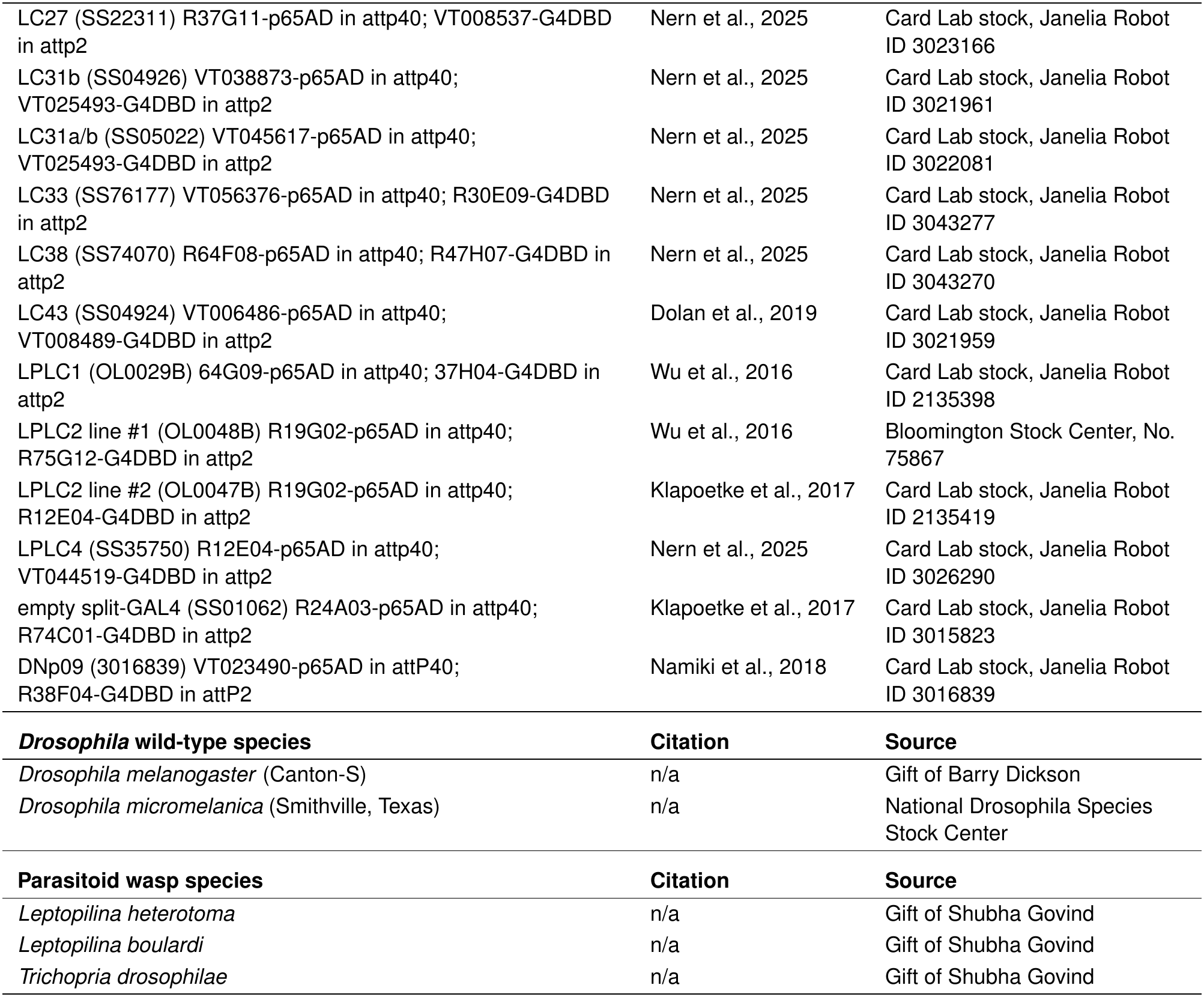
Insect stocks and sources.

| **Transgenic *Drosophila melanogaster* genotype** | **Citation** | **Source** |
|---|---|---|
| *norpA*<sup>P24</sup> | Ostroy and Pak, 1974 | Bloomington Stock Center, No. 9048 |
| UAS-TNT-E (Chr II) | Sweeney et al., 1995 | Gift of Barry Dickson |
| UAS-GCaMP8m in su(Hw)attP5 | Zhang et al., 2023 | Bloomington Stock Center, No. 605072 |
| LC4 line #1 (SS00315) R47H03-p65AD in attp40; R72E01-G4DBD in attp2 | Wu et al., 2016 | Bloomington Stock Center, No. 68259 |
| LC4 line #2 (SS00356) R47H03-p65AD in attp40; R86D05-G4DBD in attp2 | von Reyn et al., 2017 | Card Lab stock, Janelia Robot ID 3000745 |
| LC6 (OL0070B) R91B01-p65AD in attp40; R22A07-G4DBD in attp2 | Wu et al., 2016 | Bloomington Stock Center, No. 68348 |
| LC9 (SS02645) VT027704-p65AD in attp40; VT040569-G4DBD in attp2 | Wu et al., 2016 | Bloomington Stock Center, No. 68243 |
| LC10a (OL0019B) 35D04-p65AD in attp40; 22D06-G4DBD in attp2 | Wu et al., 2016 | Card Lab stock, Janelia Robot ID 2135388 |
| LC10a-d (SS02664) R35D04-p65AD in attp40; VT043656-G4DBD in attp2 | Wu et al., 2016 | Bloomington Stock Center, No. 68339 |
| LC11 line #1 (OL0015B) R22H02-p65AD in attp40; R20G06-G4DBD in attp2 | Wu et al., 2016 | Bloomington Stock Center, No. 68362 |
| LC11 line #2 (OL0043B) R55C04-p65AD in attp40; R14C08-G4DBD in attp2 | Nern et al., 2025 | Card Lab stock, Janelia Robot ID 2135415 |
| LC12 (OL0007B) R35D04-p65AD in attp40; R65B05-G4DBD in attp2 | Wu et al., 2016 | Card Lab stock, Janelia Robot ID 2135376 |
| LC13 (OL0027B) R14A11-p65AD in attp40; R50C10-G4DBD in attp2 | Wu et al., 2016 | Bloomington Stock Center, No. 68257 |
| LC14 (OL0037B) R87D07-p65AD in attp40; R59B03-G4DBD in attp2 | Nern et al., 2025 | Card Lab stock, Janelia Robot ID 2135406 |
| LC15 line #1 (OL0003B) R24A02-p65AD in attp40; R26A03-G4DBD in attp2 | Wu et al., 2016 | Bloomington Stock Center, No. 68338 |
| LC15 line #2 (OL0004B) 24A02-p65AD in attp40; 91B01-G4DBD in attp2 | Wu et al., 2016 | Card Lab stock, Janelia Robot ID 2135373 |
| LC16 line #1 (OL0092C) R92B02-p65AD in VK00027; R38E08-G4DBD in attp2 | Wu et al., 2016 | Bloomington Stock Center, No. 68376 |
| LC16 line #2 (OL0017B) 82D11-p65AD in attp40; 54A05-G4DBD in attp2 | Wu et al., 2016 | Card Lab stock, Janelia Robot ID 2135386 |
| LC17 (OL0006B) R21D03-p65AD in attp40; R24A03-G4DBD in attp2 | Nern et al., 2025 | Card Lab stock, Janelia Robot ID 2135375 |
| LC18 (OL0010B) R92B11-p65AD in attp40; R82D11-G4DBD in attp2 | Wu et al., 2016 | Bloomington Stock Center, No. 68358 |
| LC20 (SS00343) R17A04-p65AD in VK00027; R35B06-G4DBD in attp2 | Wu et al., 2016 | Bloomington Stock Center, No. 68260 |
| LC21 line #1 (SS17689) R50C03-p65AD in attp40; R41C05-G4DBD in attp2 | Klapoetke et al., 2022 | Card Lab stock, Janelia Robot ID 3022209 |
| LC21 line #2 (SS05037) R85F11-p65AD in attp40; R45F08-G4DBD in attp2 | Klapoetke et al., 2022 | Card Lab stock, Janelia Robot ID 3022096 |
| LC22 (OL0001B) R64G10-p65AD in attp40; R35B06-G4DBD in attp2 | Wu et al., 2016 | Bloomington Stock Center, No. 68357 |
| LC24 (SS02638) VT038216-p65AD in attp40; VT026477-G4DBD in attp2 | Wu et al., 2016 | Bloomington Stock Center, No. 68340 |
| LC25 line #1 (SS25130) R37D11-p65AD in attp40; VT002021-G4DBD in attp2 | Nern et al., 2025 | Card Lab stock, Janelia Robot ID 3024236 |
| LC25 line #2 (SS02650) VT009792-p65AD in attp40; VT002021-G4DBD in attp2 | Wu et al., 2016 | Bloomington Stock Center, No. 68341 |
| LC26 (SS02445) VT007747-p65AD in attp40; R85H06-G4DBD in attp2 | Wu et al., 2016 | Bloomington Stock Center, No. 68333 |
| LC27 (SS22311) R37G11-p65AD in attp40; VT008537-G4DBD in attp2 | Nern et al., 2025 | Card Lab stock, Janelia Robot ID 3023166 |
| LC31b (SS04926) VT038873-p65AD in attp40; VT025493-G4DBD in attp2 | Nern et al., 2025 | Card Lab stock, Janelia Robot ID 3021961 |
| LC31a/b (SS05022) VT045617-p65AD in attp40; VT025493-G4DBD in attp2 | Nern et al., 2025 | Card Lab stock, Janelia Robot ID 3022081 |
| LC33 (SS76177) VT056376-p65AD in attp40; R30E09-G4DBD in attp2 | Nern et al., 2025 | Card Lab stock, Janelia Robot ID 3043277 |
| LC38 (SS74070) R64F08-p65AD in attp40; R47H07-G4DBD in attp2 | Nern et al., 2025 | Card Lab stock, Janelia Robot ID 3043270 |
| LC43 (SS04924) VT006486-p65AD in attp40; VT008489-G4DBD in attp2 | Dolan et al., 2019 | Card Lab stock, Janelia Robot ID 3021959 |
| LPLC1 (OL0029B) 64G09-p65AD in attp40; 37H04-G4DBD in attp2 | Wu et al., 2016 | Card Lab stock, Janelia Robot ID 2135398 |
| LPLC2 line #1 (OL0048B) R19G02-p65AD in attp40; R75G12-G4DBD in attp2 | Wu et al., 2016 | Bloomington Stock Center, No. 75867 |
| LPLC2 line #2 (OL0047B) R19G02-p65AD in attp40; R12E04-G4DBD in attp2 | Klapoetke et al., 2017 | Card Lab stock, Janelia Robot ID 2135419 |
| LPLC4 (SS35750) R12E04-p65AD in attp40; VT044519-G4DBD in attp2 | Nern et al., 2025 | Card Lab stock, Janelia Robot ID 3026290 |
| empty split-GAL4 (SS01062) R24A03-p65AD in attp40; R74C01-G4DBD in attp2 | Klapoetke et al., 2017 | Card Lab stock, Janelia Robot ID 3015823 |
| DNp09 (3016839) VT023490-p65AD in attP40; R38F04-G4DBD in attP2 | Namiki et al., 2018 | Card Lab stock, Janelia Robot ID 3016839 |
| ***Drosophila* wild-type species** | **Citation** | **Source** |
| *Drosophila melanogaster* (Canton-S) | n/a | Gift of Barry Dickson |
| *Drosophila micromelanica* (Smithville, Texas) | n/a | National Drosophila Species Stock Center |
| **Parasitoid wasp species** | **Citation** | **Source** |
| *Leptopilina heterotoma* | n/a | Gift of Shubha Govind |
| *Leptopilina boulardi* | n/a | Gift of Shubha Govind |
| *Trichopria drosophilae* | n/a | Gift of Shubha Govind |

